# Tension-driven multi-scale self-organisation in human iPSC-derived muscle fibers

**DOI:** 10.1101/2021.10.24.465614

**Authors:** Qiyan Mao, Achyuth Acharya, Alejandra Rodríguez-delaRosa, Fabio Marchiano, Benoit Dehapiot, Ziad Al Tanoury, Jyoti Rao, Margarete Díaz-Cuadros, Arian Mansur, Erica Wagner, Claire Chardes, Vandana A. Gupta, Pierre-François Lenne, Bianca H. Habermann, Olivier Theodoly, Olivier Pourquie, Frank Schnorrer

**Affiliations:** Aix Marseille University, CNRS, IBDM, Turing Centre for Living Systems, Marseille, France; Department of Pathology, Brigham and Women’s Hospital, Boston, MA, USA; Department of Genetics, Harvard Medical School, Boston, MA, USA; Harvard Stem Cell Institute, Boston, MA, USA; Division of Genetics, Department of Medicine, Brigham and Women’s Hospital, Harvard Medical School, Boston, MA, USA; Aix Marseille University, CNRS, LAI, Turing Centre for Living Systems, Marseille, France

**Keywords:** muscle, myofibril, sarcomere, human induced pluripotent stem cells, self-organisation, mechanical tension

## Abstract

Human muscle is a hierarchically organised tissue with its contractile cells called myofibers packed into large myofiber bundles. Each myofiber contains periodic myofibrils built by hundreds of contractile sarcomeres that generate large mechanical forces. To better understand the mechanisms that coordinate human muscle morphogenesis from tissue to molecular scales, we adopted a simple *in vitro* system using induced pluripotent stem cell-derived human myogenic precursors. When grown on an unrestricted two-dimensional substrate, developing myofibers spontaneously align and self-organise into higher-order myofiber bundles, which grow and consolidate to stable sizes. Following a transcriptional boost of sarcomeric components, myofibrils assemble into chains of periodic sarcomeres that emerge across the entire myofiber. By directly probing tension we found that tension build-up precedes sarcomere assembly and increases within each assembling myofibril. Furthermore, we found that myofiber ends stably attach to other myofibers using integrin-based attachments and thus myofiber bundling coincides with stable myofiber bundle attachment *in vitro*. A failure in stable myofiber attachment results in a collapse of the myofibrils. Overall, our results strongly suggest that mechanical tension across sarcomeric components as well as between differentiating myofibers is key to coordinate the multi-scale self-organisation of muscle morphogenesis.

## Introduction

Human skeletal muscles display a very hierarchical and stereotypical architecture. At its distal ends, each muscle is mechanically connected to different skeletal elements, which are moved by the contracting muscle to achieve locomotion. Within each muscle, contractile muscle cells called myofibers are organised into myofiber bundles, several of which are combined to form one human muscle (Hill and Olson, 2012). Within each muscle fiber, the contractile myofibrils are laterally aligned displaying the well-appreciated cross-striated morphology of skeletal muscle fibers (Huxley and Niedergerke, 1954; Huxley and Hanson, 1954). Each myofibril consists of a long chain of contractile machines called sarcomeres, which extend across the entire length of the muscle fiber to mechanically connect the attached skeletal elements (Schiaffino and Reggiani, 2011). As such, coordinated contraction of all sarcomeres shortens all myofibrils and thus shortens each muscle fiber. This results in force production powering the desired body movement (Lemke and Schnorrer, 2017; Lindstedt, 2016).

Sarcomere architecture is highly regular. The plus ends of parallel actin filaments are cross-linked at the Z-discs bordering each sarcomere, while their minus ends point centrally where they overlap with large bipolar myosin filaments. The latter are cross-linked in the middle of the sarcomere building the M-band (Dasbiswas et al., 2018; Lange et al., 2006). Both types of filaments are stably linked together by the gigantic elastic protein titin, whose N-terminus is bound to *α*-actinin at the Z-disc, while its C-terminus is embedded into the M-band (Lange et al., 2006; Linke, 2018; Tskhovrebova and Trinick, 2003). This results in a stereotypic sarcomere length, depending on muscle type ranging from two to three micrometres in relaxed human muscle *in vivo* (Ehler and Gautel, 2008; Llewellyn et al., 2008; Tskhovrebova and Trinick, 2003).

As most mature human muscles are several centimetres long, many thousand sarcomeres need to be concatenated into each myofibril to mechanically connect both myofiber ends (Lemke and Schnorrer, 2017). Assembling the correct number of sarcomeres will result in the optimal amount of passive tension present in the relaxed muscle. This is a prerequisite for effective active muscle force production during sarcomere contraction, powered by myosin forces pulling on actin (Dasbiswas et al., 2018; Linke, 2018). Hence, the complex hierarchical muscle tissue operates under continuous mechanical tension.

Optimal tension levels are not only important in the mature muscle, but also essential during its development. Insights from *in vivo* imaging of myofibrillogenesis during *Drosophila* flight muscle development sparked the tension-driven self-organisation hypothesis of myofibrillogenesis (Weitkunat et al., 2014): this hypothesis suggests that myotubes first attach to tendons. After successful attachment, they build up mechanical tension along the long axis of the muscle fiber. This tension build-up triggers the simultaneous assembly of long periodic myofibrils that extend through the entire fiber and thus mechanically connect the two attached exoskeletal elements of the fly (Weitkunat et al., 2014; 2017). Thus, tension is key to coordinate myofibril self-organisation over long distances in insect muscles.

However, *Drosophila* muscles are simple, single myofiber muscles that do not form higher order myofiber bundles. Hence, it is a limited model for understanding how myofiber attachment, myofiber bundling and sarcomere assembly are coordinated during human muscle tissue morphogenesis. Furthermore, human muscle tissue not only connects to tendon and cartilage cells, which are the source of various guidance cues *in vivo* (Schweitzer et al., 2010), but also contains a complex muscle-associated connective tissue that may further instruct muscle development both mechanically and chemically (Helmbacher and Stricker, 2020). How precisely this complex interplay of cell types may coordinate muscle morphogenesis at different scales is not well understood.

Here we set out to test the self-organisation capacity of human muscle tissue *in vitro* when developing on a substrate lacking any localised guidance, attachment or other mechanical cues. We employed a recently developed human induced pluripotent stem cell (iPSC) differentiation protocol that can efficiently generate myogenic precursors which further differentiate robustly into striated skeletal muscle fibers *in vitro* (Chal et al., 2015; 2016; Tanoury et al., 2021; Yan et al., 2021). This method allowed us to obtain large quantities of isogenic human myogenic precursors of defined developmental history, in contrast to primary myoblast cultures from donors. Furthermore, CRISPR-based genome engineering enabled us to tag sarcomeric protein components of choice to follow their live dynamics *in vitro* (Sharma et al., 2018; Tanoury et al., 2021).

Using two-dimensional cultures of human iPSC-derived myogenic precursors, we found that despite the absence of attachment cues, myofibers elongate to self-restricted stable lengths. Concomitantly, they self-organise into large myofiber bundles that attach to shared attachment foci at both ends. As myofibers approach stable lengths, their sarcomeres emerge simultaneously over broad regions, suggesting that myofibril and myofiber morphogenesis are coordinated from tissue to molecular scales. We identified mechanical tension as a likely coordination mechanism as tension precedes sarcomere formation and further increases when myofibrils mature. Consequently, integrin-based adhesions between the bundled myofiber ends support stable myofiber attachment. Since tension can be transmitted across tissue, cell and molecular scales, we hypothesize that tension is key to coordinate muscle morphogenesis across scales.

## Results

### Human iPSC-derived myofibers follow a biphasic differentiation process

In order to study human muscle fiber morphogenesis on cellular and subcellular scales, we adapted a 2D *in vitro* culture system. We generated human iPSC-derived myogenic precursors by differentiating NCRM1 iPS cells for 20-30 days according to [18]. These cultures were then dissociated and filtered through 70 µm and then 40 µm cell filters to obtain single-cell suspensions. These were replated in SKGM-2 medium for 1 to 2 days to generate cultures enriched in myogenic precursors prior to being frozen (Chal et al., 2016; Tanoury et al., 2021). Frozen myogenic precursors were thawed, seeded on Matrigel-coated ibidi® glass-bottom dishes and grown in proliferation medium (SKGM-2) for 2-3 days before being switched to KCTiP differentiation medium (see materials and methods). On day 1 in KCTiP differentiation medium, the myogenic precursors initially resemble fibroblasts by shape (Figure 1A). These myogenic precursors are able to fuse to other myogenic precursors or elongated myotubes to form multi-nucleated myotubes (Figure 1 – movie 1, Figure 1 – figure supplement 1A).

**Figure 1.**
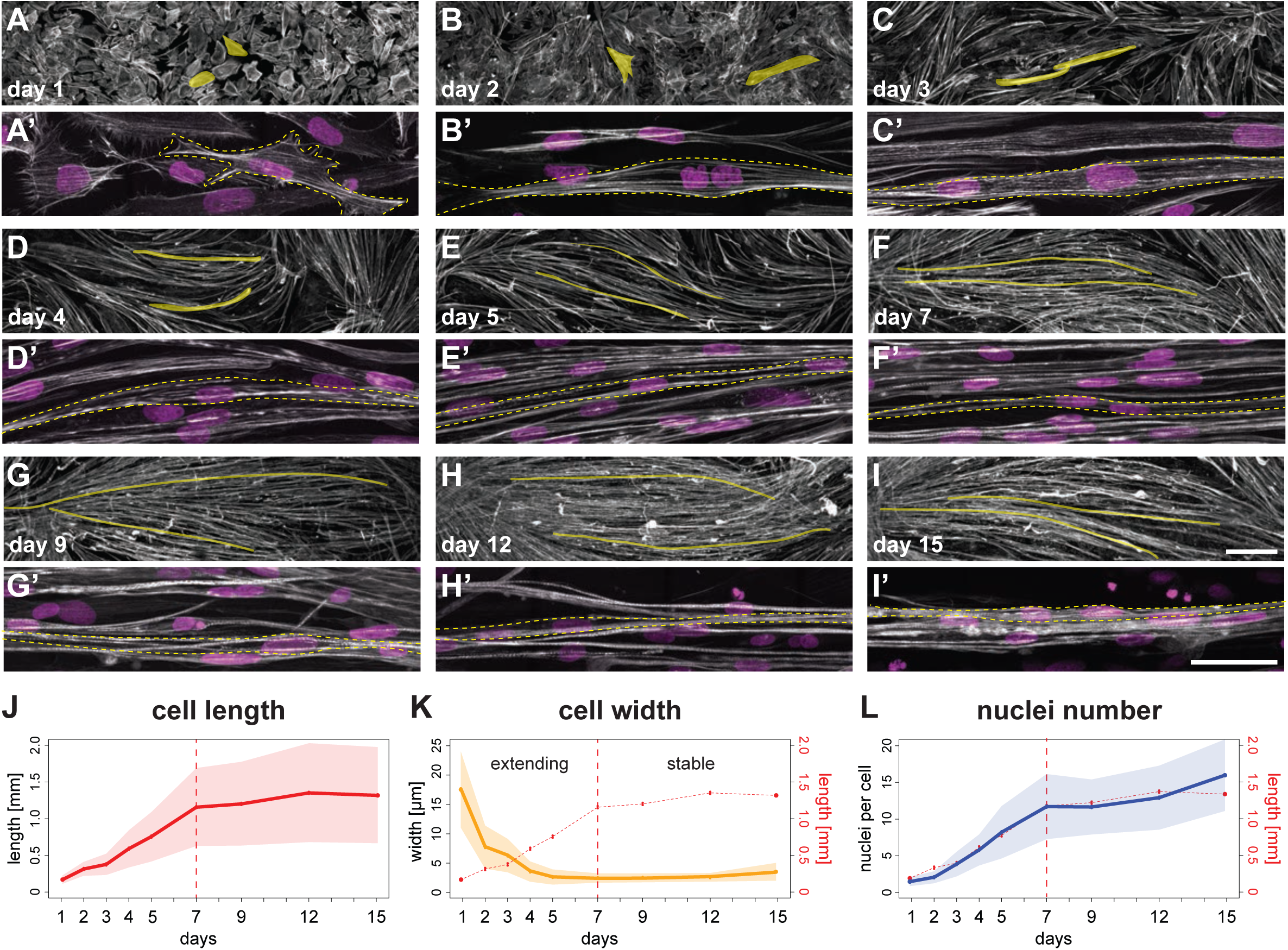
A time course of human iPSC-derived myofibers in a minimalist 2D culture. **A-I)** 10× views of phalloidin staining of human iPSC-derived myofibers. Yellow shades highlight two randomly selected myocytes, myotubes or myofibers. **A’-I’)** 100× views of individual myocytes, myotubes or myofibers. Rhodamine-phalloidin in grey, DAPI in magenta. Yellow dashed lines highlight individual myocytes/myotubes/myofibers. **J)** Length of myocytes/myotubes/myofibers over time (n = 35, 42, 4695, 3765, 4191, 2439, 2061, 1880, 1883 myofibers per reported day, respectively). Dots and solid line show the mean of each day, while shaded area shows the standard deviation. Vertical dashed line shows the transition point at day 7 from the elongation phase to the stable phase. **K)** Width of myocytes/myotubes/myofibers over time (n = 45, 50, 111, 191, 175, 297, 340, 315, 201 myofibers per reported day, respectively). L) Number of nuclei per myofiber over time (n = 35, 42, 76, 81, 85, 77, 62, 74, 66 myofibers per day). In K) and L), dots, shaded area and solid line follow convention in J), with graph in J) is reproduced in red. Scale bars: 250 µm in A-I), 50 µm in A’-I’). **[Related to Figure 1 – figure supplement 1, and Figure 1 – video 1. Source data available for Figure 1J-L.]**

Concomitant with myogenic precursors fusion, myotubes rapidly elongate, while their width decreases during the first week (Figure 1A-F, A’-F’, J, K, Figure 1 – movie 1, Figure 1 – figure supplement 1B), suggesting that they are actively extending to increase their aspect ratio. The average length of a day 7 myotube/myofiber is about 1.2 mm, while being only 3 - 5 µm in width in our culture conditions. During the second week of culture, average myofiber length stabilises at about 1.2 - 1.3 mm (Figure 1G, K), while fusion continues, reaching an average of 15 nuclei per myofiber at day 15 (Figure 1L). This happens despite that fact that our culture dishes have an average dimension of 6 mm, suggesting that a self-limiting mechanism may prevent further elongation of the myofibers. We conclude that on our unpatterned 2D substrates, differentiating myotubes/myofibers follow a biphasic behaviour of elongation until day 7, followed by a stabilisation in length until day 15.

### Myofibers self-organise into bundles that merge to stable sizes

To better understand how myofibers reach stable lengths at day 7, we investigated their spatial organisation with respect to neighbouring myofibers. We found that from day 3 and 4, myotubes and myofibers generate swirly patterns that enlarge over time (Figure 2A-F), indicating that new myofibers are aligning with growing myofiber bundles despite the lack of predefined orientation cues. This suggests a self-organisation mechanism during myofiber bundling.

**Figure 2.**
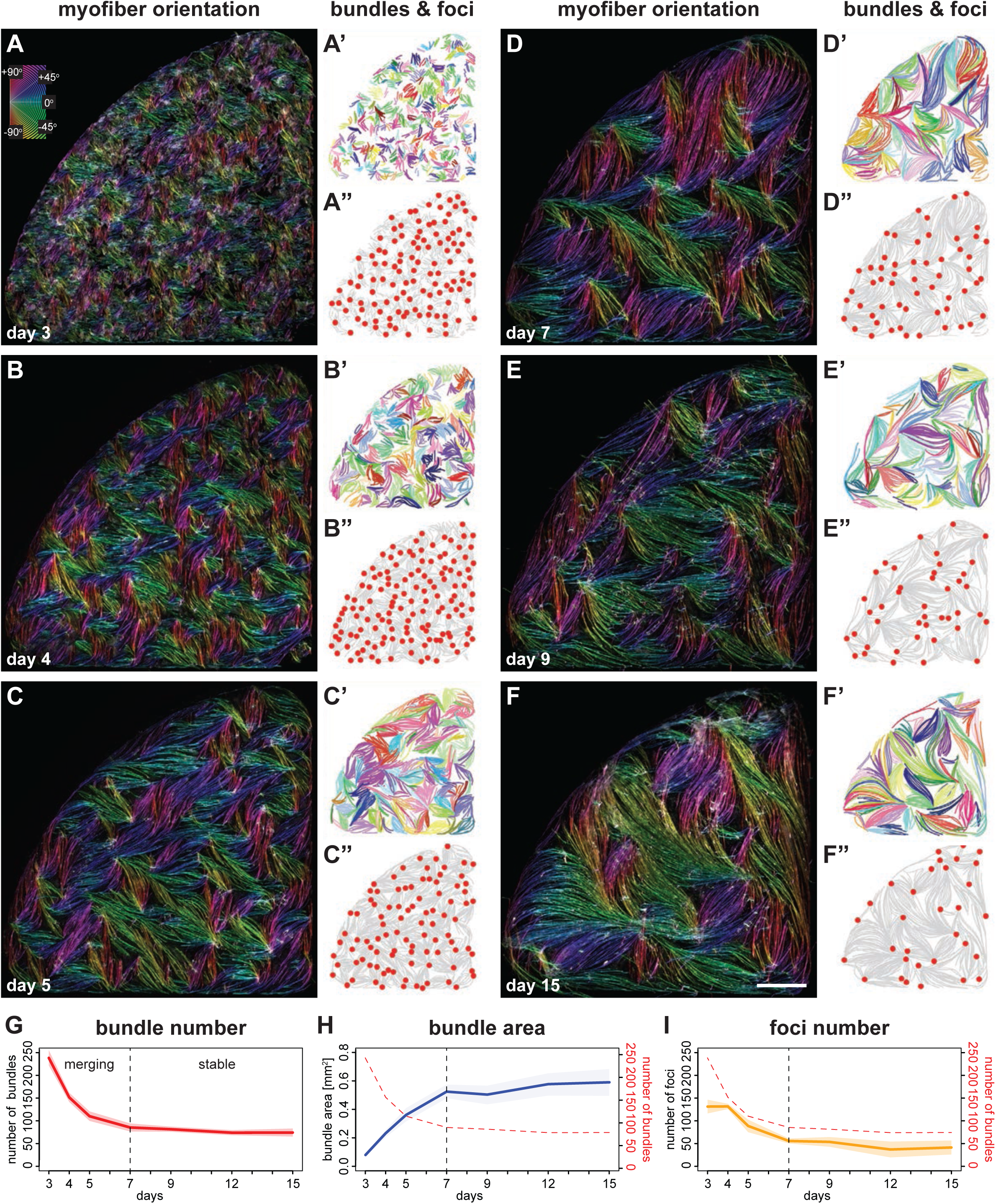
Myofibers self-organise into bundles that consolidate to stable sizes. **A-F)** Rhodamine-phalloidin staining pseudo-coloured according to local fiber orientation by OrientationJ. Inset shows a colour wheel used by OrientationJ. **A’-F’)** automatically segmented myofiber bundles from manually traced myofibers, colour-coded by bundle identity. **A”-F”)** manually traced myofiber bundle end foci (red dots) super imposed on traced myofibers (grey lines reproduced from A’-F’). **G)** Number of myofiber bundles over time (n = 7 cultures per stage). Solid line shows the mean of each day, while shaded area shows the standard deviation. Vertical dashed line shows the transition point at day 7 from a merging to a stable phase. **H)** Area of myofiber bundles over time (n = 7 cultures per stage). **I)** Number of myofiber bundle end foci over time (n = 3 cultures per stage). Myofiber bundle number from G is overlayed in red. Scale bar: 1mm. **[Related to Figure 2 – figure supplement 1, figure supplement 2, and Figure 2 – video 1 . Source data available for Figure 2G-I.]**

To quantitatively analyse the alignment of myotubes and myofibers with their neighbours, we first displayed the local fiber orientation as vector fields for the central region of the culture (Figure 2 – figure supplement 1, see methods). At day 3 we observed small swirly patterns in vector fields (Figure 2 – figure supplement 1A), indicative of short-range alignments of polarity. These aligned regions enlarge over time (Figure 2 – figure supplement 1A-F), which is supported by an increasing average nematic length during differentiation (Figure 2 – figure supplement 1G, H; nematic length was defined as the intersection of the initial linear decay of the spatial correlation function with the *x* axis, see methods). To visualise the local discontinuities of alignment, which often locate at the end of myofiber bundles, we calculated the nematic order for each vector grid (Figure 2 – figure supplement 1A’-F’; nematic order is a parameter describing the alignment of local orientation of the vector field at a given position, see methods). This defined regions of disorder in the cultures with a nematic order below a threshold of 0.5. The fraction of disordered regions decreases over time (Figure 2 – figure supplement 1I), further supporting an expansion of aligned regions and a reduction in number of myofiber bundle ends.

In order to better understand the development of individual myofiber bundles in the entire cultures, we manually traced a total of 34,962 individual myotubes or myofibers and applied an automated Hausdorff distance clustering algorithm to objectively define distinct individual myofiber bundles (Figure 2A, see methods). This analysis revealed that the number of bundles decreases within four days from about 240 bundles (in each 0.35 cm^2^ well) at day 3 to about 75 bundles at day 7 (Figure 2A’-F’, G). In turn, the average area occupied by each bundle increases about 6-fold (Figure 2H). This strongly suggests that concomitant with myofiber elongation (see Figure 1), several smaller myofiber bundles merge into larger bundles.

Furthermore, myofiber ends present in the same bundle tend to point towards defined foci, which may serve as mechanical attachment sites to the surrounding matrix. In line with the reduction of bundle number over time, the number of these foci also decreases with similar temporal dynamics from about 140 foci at day 3 to about 50 foci at day 7 (Figure 2A’’-F’’, I). This shows that these foci are dynamic and merge before day 7, suggesting that the myofiber bundles are not yet stably attached at their respective ends.

Next, we wanted to directly document the live dynamics of myofiber bundle and foci merging. Using a smaller culture format, we recorded the entire cultures by live phase-contrast imaging. Thus, we followed individual myofiber bundles over four days during differentiation and identified several examples in which one bundle elongates its extremity along a neighbouring bundle and finally merges with the neighbouring bundle to form shared attachment foci (Figure 2 – figure supplement 2, Figure 2 – video 1). This suggests that neighbouring myofiber bundles can offer orientation cues to nearby bundles during the dynamic bundle merging phase between day 3 to day 7 of the culture. This revealed that myofiber bundles can self-organise, potentially using mechanical orientation cues presented by their neighbours.

We noticed that myofiber bundles cease to merge after day 7 in our cultures (Figure 2A-G), coinciding with the end of myofiber extension (Figure 1J). Day 7 also demarcates a turning point for bundle area growth and foci fusion, both of which remain largely stable after day 7 (Figure 2H, I). We conclude that myotubes and myofibers show a remarkable biphasic differentiation behaviour *in vitro* that is coordinated between cell and tissue scales. We hypothesized that the biphasic growth of myofibers and myofiber bundles may be caused by a switch in their mechanical status around day 7. This switch may further result in molecular changes in myofibers that enable myofibril and sarcomere formation.

### A transcriptional switch to enable myofibril and sarcomere assembly

As a first step to investigate the molecular changes at day 7 of differentiation, we analysed a previously generated transcriptomics dataset comparing similar proliferative cultures (using the identical SKGM-2 culture conditions) with day 7 and day 15 of myogenic differentiation (using the identical KCTiP differentiation conditions) (Tanoury et al., 2021). After verifying the quality of the transcriptomics data set with principal components analysis (Figure 3 – figure supplement 1, Supplementary Table 1) we performed Mfuzz clustering (Kumar and Futschik, 2007), choosing 10 clusters as optimal for our data set (see methods). Mfuzz clustering assigned all expressed genes to one of the 10 clusters which exhibit different expression dynamics (Figure 3A, Supplementary Table 1, see methods). In search for clusters that may reflect the temporal dynamics of the biphasic myofiber extension behaviour we found that three gene clusters are upregulated at day 7 of differentiation, while reaching a plateau or decreasing again at day 15 (clusters 1, 2, 7). Notably two of these clusters are enriched for muscle specific GO-terms (Figure 3A, B). In particular, cluster 1 contains many genes coding for sarcomeric protein components, whose expression shows a strong upregulation at day 7 and then plateaus until day 15 (Figure 3A, B, Supplementary Table 1). Furthermore, cluster 5, which is also strongly upregulated at day 7 and remains high at day 15 is also enriched for striated muscle contraction GO-terms (Figure 3A, B, Supplementary Table 1). In contrast, components of the mitochondrial respiratory chain are only upregulated after day 7 and hence enriched in cluster 3 and 4 (Figure 3A, B, Supplementary Table 1). This indicated that the sarcomere components are selectively upregulated at day 7, coinciding with the observed myofiber bundle consolidation.

**Figure 3.**
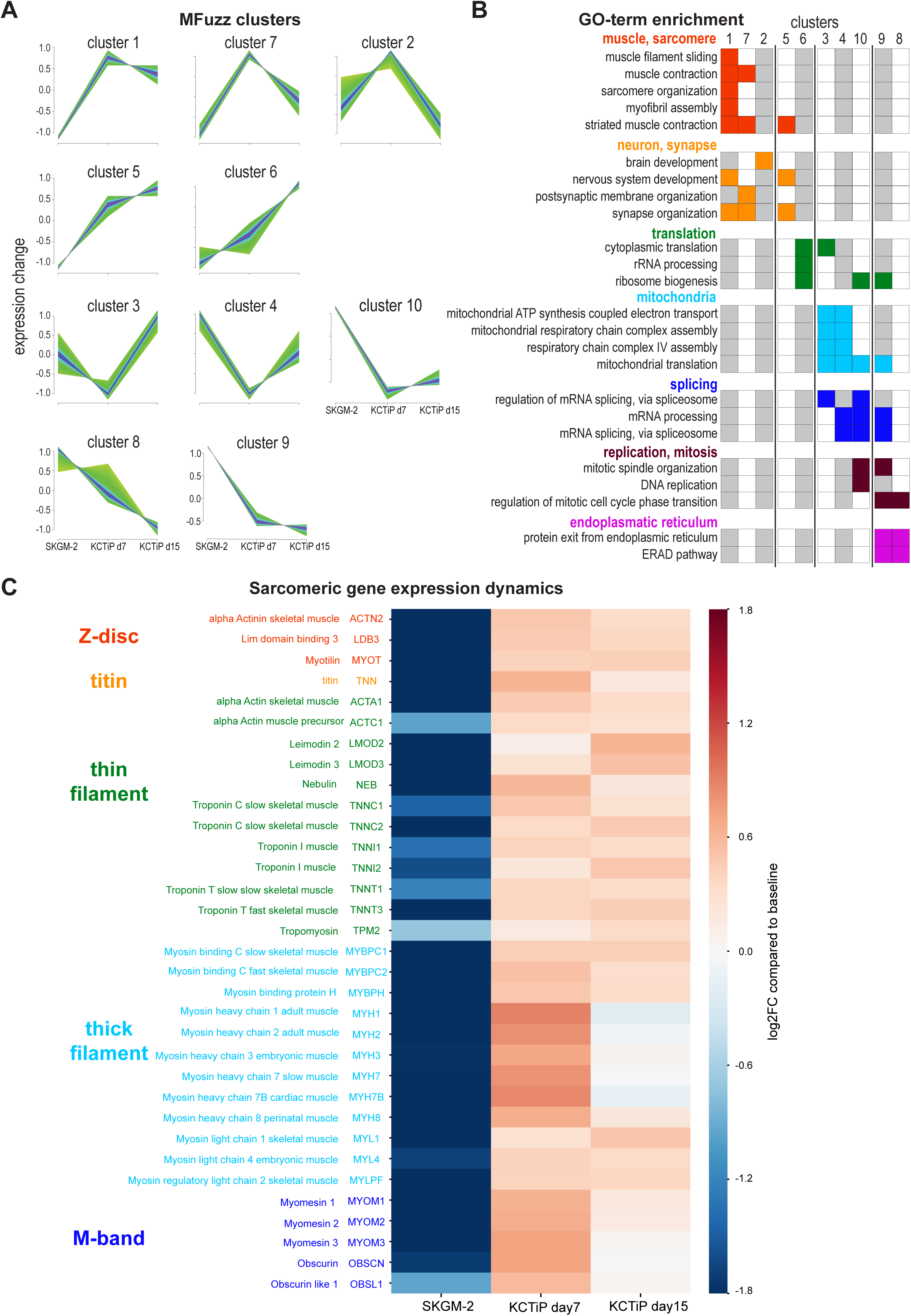
A transcriptional switch boosts sarcomeric gene expression. **A)** Mfuzz clustering defined 10 distinct gene clusters with shared expression dynamics over the time course. **B)** Selected enriched GO-terms for the respective clusters are displayed, a full list of GO-term enrichments is presented in Supplementary Table 1. Note the two muscle-enriched Mfuzz clusters displaying a stark expression increase from undifferentiated to day 7. **C)** Heatmap showing expression dynamics of various key sarcomeric genes comparing undifferentiated, day 7 and day 15 myogenic cultures to baseline. Note the stark expression increase of all components until day 7. **[Related to Figure 3 – figure supplement 1.]**

Sarcomere assembly requires the concerted action of a large number of sarcomeric components (Lange et al., 2006). Closer inspection of the highly expressed and hence relevant genes coding for sarcomere components revealed that indeed all of them are strongly upregulated, many of them more than 10-fold at day 7 compared to proliferating cultures (Figure 3C, Supplementary Table 1). This includes thin filament components actin A1, nebulin, leimodin 2, 3 and various troponin subunits, as well as thick filament components myosin heavy chain 1, 2, 3, 7, 8, myosin light chain 1, 3 and myosin binding protein C1, 2, and finally the titin protein, the Z-disc components *α*-actinin 2, the ZASP homolog LDB3 and the M-band core members obscurin and myomesin 1, 2 and 3 (Figure 3C). Many of these sarcomeric components are not further upregulated comparing day 7 to day 15 and hence are members of clusters 1 and 7, while expression of some does slightly increase further and hence these are members of cluster 5 (Figure 3A, C, Supplementary Table 1). This suggests that the transcriptional switch inducing expression of the sarcomeric components at day 7 correlates well with the biphasic morphogenesis behaviour of myofibers reaching a stable length after day 7.

### Periodic sarcomeres emerge as myofibers approach stable lengths

The emergence of stable attachment foci during myotube differentiation and the strong induction of sarcomere gene expression at day 7 motivated a careful analysis of myofibrillogenesis and sarcomerogenesis in the differentiating cultures. We immuno-stained the cultures to visualise actin (with phalloidin), muscle myosin (with myosin heavy chain antibody recognising all MyHC isoforms (Webster et al., 1988)) and titin (with a N-terminal titin epitope antibody located at the sarcomeric Z-disc (Mayans et al., 1998)) and manually segmented 810 myotubes or myofibers. Each fiber was then assigned to one of four morphologically distinct categories (Figure 4A): first, ‘myotubes’ display very low levels of titin in a salt-and-pepper pattern, muscle myosin levels are also low and actin does not yet form long continuous myofibrils. The lack of myofibrils is substantiated by the absence of periodic myosin or titin patterns on actin filaments (Figure 4A, B). Thus, myotubes do not yet contain myofibrils. Second, ‘immature myofibers’ display higher levels of titin and myosin, both of which are recruited onto continuous actin structures, which do not yet display an obvious periodicity (Figure 4A, B). Thus, immature myofibers contain immature myofibrils that are not yet periodic. Third, ‘transitional myofibers’ show high titin and myosin levels, both of which display a periodic pattern on long continuous periodic actin structures (Figure 4A, B). Hence, transitional myofibers contain long periodic myofibrils built from chains of immature sarcomeres. Fourth, ‘mature myofibers’ display strongly periodic patterns of titin, myosin and actin across the entire width of the myofiber (Figure 4A, B). Thus, mature myofibrils in mature myofibers have either aligned or laterally expanded to form large Z-discs. Generally, the patterns appeared homogenous throughout most of the length of the myotube or myofiber; hence, we were able to categorise each of them into a single morphological category.

**Figure 4.**
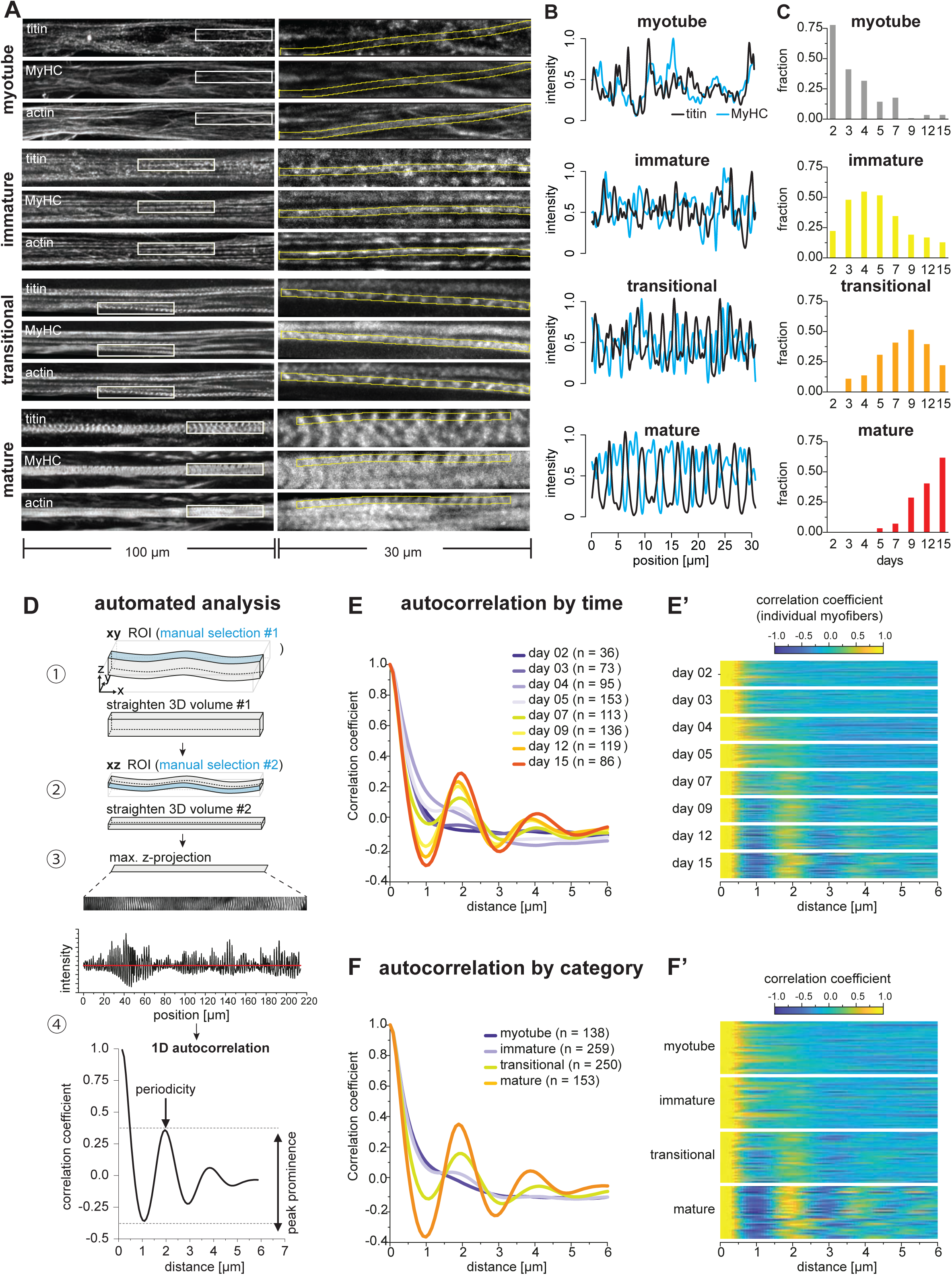
Sarcomeres emerge as myofibers approach stable lengths. **A)** Multi-channel fluorescent images of four morphological categories (myotubes, immature, transitional and mature myofibers). Left column shows a view of 100 µm wide, right column a zoomed view of 30 µm boxed area. Immunofluorescence against titin N-terminus, muscle myosin heavy chain (MyHC), and phalloidin staining of actin are shown. **B)** Normalised 1D fluorescent intensities for titin (black) and myosin (blue) from ROIs in right column in A. Note the emerging periodicity in transitional fibers. **C)** Distribution of each morphological category during the time course. **D)** Semi-automatic 1D autocorrelation analysis pipeline. Step ①: Straightening manual ROI in xy dimension. Step ②: Straightening ROI from Step ① in xz dimension. Step ③: Generating 1D intensity profile along myofiber. Step ④: autocorrelation analysis (ACF) with 1D intensity profile from Step ③. Presence of a secondary peak indicates the existence of periodic signal, with peak position indicating period length and peak prominence reflecting period regularity. **E)** Average ACF for each time point. **E’)** Randomly selected ACFs for each time point (n=36 each). **F)** Average ACF for each morphological category. **F’)** Randomly selected ACFs for each category (n=36 each). **[Source data available for Figure 4C.]**

Having these four categories in hand, we classified all the 810 traced myotubes and myofibers across the entire time course to gain insights into the evolution of myofiber morphology at the population level (Figure 4C). Myotubes and immature myofibers are largely present before day 7 in the differentiating cultures, whereas transitional and mature myofibers dominate after day 7 (Figure 4C). This fits well with the observed biphasic behaviour described above and suggests that periodic sarcomeres largely assemble around day 7, the time when myofibers become stably attached.

In order to analyse the time course of sarcomere assembly in an unbiased manner, we developed a semi-automatic image analysis pipeline to quantitate periodic titin signal intensity profiles in a 3D volume extending along a length of up to 200 µm using autocorrelation function analysis (ACF) (Figure 4D, see methods). After straightening the segmented myofibril to obtain a 1D titin intensity profile, we applied an autocorrelation function and monitored secondary ACF peaks position and prominence as a read out of the regularity of the pattern (Figure 4D). We applied this pipeline to all myotubes and myofibers fixed at different time points in the differentiation time course. When averaging all ACFs for each time point, we found a secondary peak first emerging at days 5 and 7, with a period length of about 2 µm (Figure 4E). This sarcomere length fits well with the length of human fetal muscle sarcomeres (Racca et al., 2013). The peak prominence becomes higher at day 9, 12 and 15 (Figure 4E), demonstrating that the periodicity and thus the sarcomeres mature at later stages. This is also visible from individual ACFs in randomly selected myotubes or myofibers from each time point (Figure 4E’). Finally, we sorted the ACF based on our manually assigned morphological categories and confirmed that immature myofibers do not yet display any obvious periodicity. Periodicity is however clearly detected in transitional and mature myofibers (Figure 4F, F’). We conclude that periodic sarcomeres assemble in transitional myofibers around days 5 to 7. This coincides with the time when myofibers have formed stable bundles with stable attachment foci (Figures 1 and 2). This correlation of morphological transitions across molecular-, cellular- and tissue-scales led us to hypothesise that mechanical signals may serve as a common denominator of morphological organisations.

### Sarcomeres emerge simultaneously over broad regions in myofibers

*Drosophila* muscles require mechanical tension to trigger the simultaneous assembly of periodic myofibrils across the entire length of the myofiber (Weitkunat et al., 2014). If mechanical tension also provides a long-range coordination signal in human myofiber cultures, then one would predict that sarcomeres may also emerge across broad regions after stable attachment.

In order to study how the first periodic myofibrils in human fibers are built, we performed local ACF analysis aiming to analyse sarcomere periodicity in 10 µm-wide windows sliding along the ‘straightened’ 1D titin intensity profiles ranging from 90 to 200 µm in length. This generated a 2D periodicity map for each myofiber, displaying the regularity of the pattern, measured by the peak prominence (Figure 5A, white line represents peak prominence), and distance between peaks (Figure 5A). To minimise artefacts generated by our straightening procedure we optimised the peak prominence threshold and allowed for short gaps (below 6 µm in length; see methods). This enabled us to quantify the periodicity of titin distribution in all our 810 myotubes and myofibers (red lines in Figure 5A). As expected from the global ACF analysis shown in Figure 4, myotubes display almost no titin periodic region above the threshold, confirming that they do not contain any myofibrils or sarcomeres (Figure 5A, B). Similarly, immature myofibers contain on average less than one periodically patterned region per 100 µm, again verifying that they do not contain periodic myofibrils yet (Figure 5A, B). Therefore, myotubes and immature myofibers have not yet assembled periodic myofibrils and hence do not yet contain sarcomeres. In contrast, our ACF analysis revealed that mature myofibers are periodically patterned across 80% of the analysed regions, showing mostly one or two large regions (Figure 5A-C). This demonstrates that mature myofibers indeed contain highly regular periodic myofibrils that span most of the myofiber.

**Figure 5.**
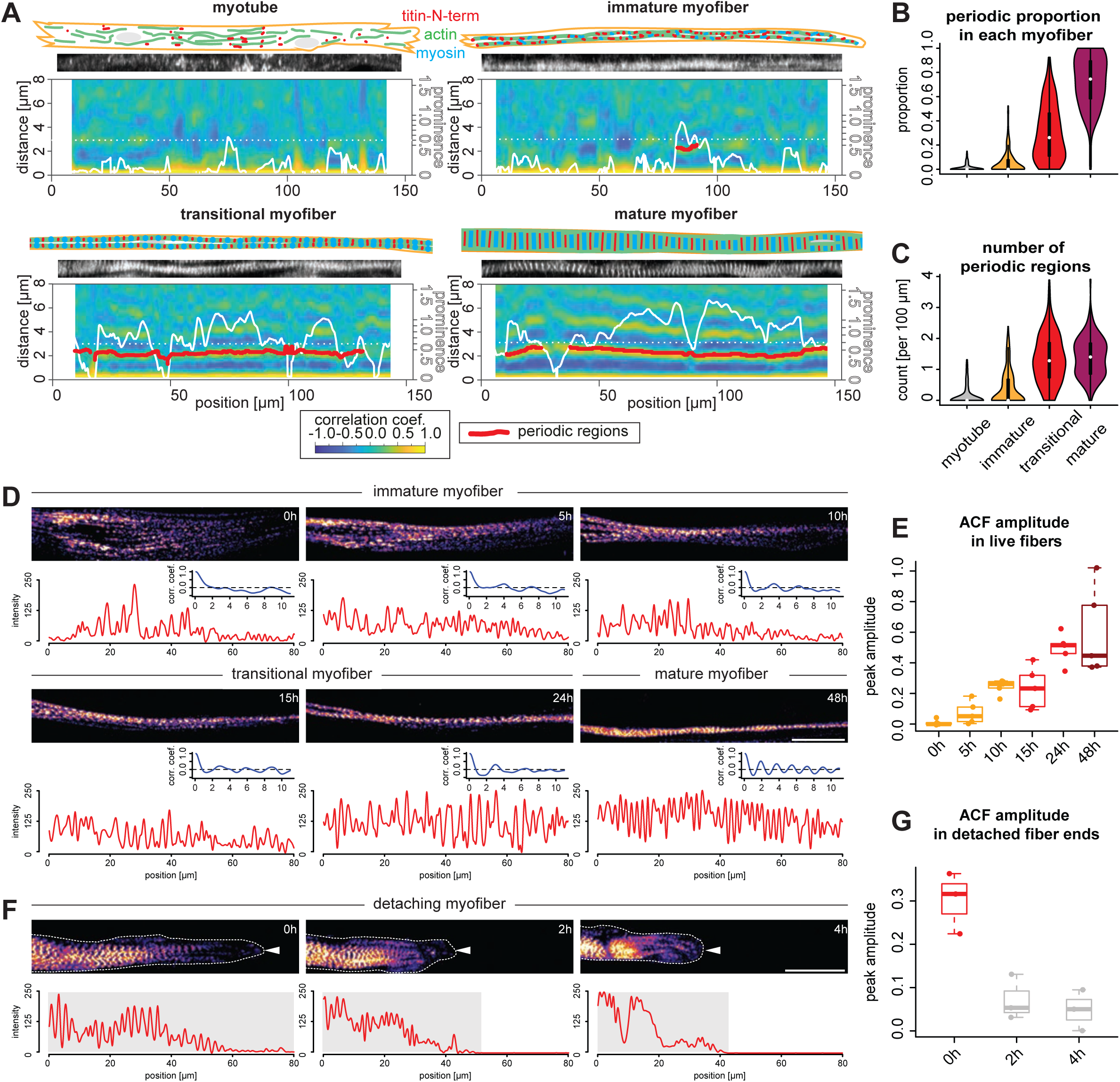
Sarcomeres emerge simultaneously over broad regions within myofibers. **A)** Local autocorrelation analysis (ACF) for each morphological category (myotubes, immature, transitional and mature myofibers). For each panel, top: schematic model of the myotube and myofiber stage with actin, myosin and titin as indicated; middle: a straightened image of titin immunofluorescence; bottom: a 2D colormap of local ACF. x-axis shows the position along the myofiber, y-axis shows the ACF distance. Superimposed white line shows prominence of the secondary peak along the myofiber. Superimposed red segments highlights secondary peak positions classified as periodic (see Methods). **B-C)** Violin plots of proportion of total length of detected periodic regions over whole lengths of myofiber (B) and number of detected periodic regions (C) for each myofiber from each category. Black bars mark the distance between 1^st^ and 3^rd^ quadrant. White dots mark the median values. **D)** Stills from a time-lapse movie showing a myofiber during myofibrillogenesis (labelled with mKate2-α-actinin2, Figure 5 – video 1). Note that the end of the movie the myofibrils displays a periodic pattern over the entire traced length, corresponding to the morphology of a mature myofiber. In each panel, the myofiber is traced to produce the intensity profile (red) and an autocorrelation function (blue) below that panel. The amplitude of the blue curve is then used to calculate values in E). **E)** Amplitudes of auto-correlation functions at different time points in myofibers undergoing myofibrillogenesis (n = 5). **F)** Stills from a time-lapse movie showing one myofiber end retracting (labelled with mKate2-α-actinin2 and outlined with white dashed line and a white arrowhead, Figure 5 – video 2). Note that the periodicity of the initially periodic myofiber collapses. In each panel, the myofiber is traced to produce the intensity profile (red) below that panel. The amplitude of the autocorrelation from the shaded region is then used to calculate values in G). **G)** Amplitudes of auto-correlation functions at different time points in collapsing myofibers (n = 3). Scale bars: 20 µm. **[Related to Figure 5 – figure supplement 1, Figure 5 – video 1 and video 2. Source data available for Figure 5B, C, E, G.].**

Together, this suggests that periodic myofibrils first assemble in transitional myofibers. This is supported by the finding that ACF analysis scores about 30% of their length as periodic (Figure 5A, B). Consistently, we find that transitional myofibers contain an average of 1.5 patterned regions per 100 µm (Figure 5C), some developed already into very long continuous regions (>50 µm, Figure 5A). This verified that transitional myofibers are the stage at which periodic sarcomeres assemble over long distances.

We next investigated the distribution of periodic regions along the length of transitional myofibers. If sarcomeres assemble simultaneously along the myofiber, our detection threshold should be reached at random positions, homogeneously distributed along the myofiber. Indeed, we find that patterned regions in transitional myofibers are homogeneously distributed throughout the entire fiber with no bias to the centre or to distal ends of the fibers (Figure 5 – figure supplement 1). Taken together, this strongly suggests that sarcomeres emerge rather simultaneously over broad regions to form long periodic myofibrils in human myofibers.

To directly visualise sarcomerogenesis in human muscle cultures live, we tagged the endogenous α-actinin2 protein with mKate2 using CRISPR-Cas9 technology (see methods). Performing fluorescent time-lapse imaging, we were able to track individual myofibers in low density cultures for up to 3 days, which enabled us to visualise myofibrillogenesis directly (Figure 5D). As observed in fixed cultures, we found that initially unpatterned immature myofibers develop periodic myofibrils over a broad region of the myofiber (Figure 5D and E, Figure 5 – video 1). This likely coincides with a strong increase in mechanical tension as we frequently found one end of the myofiber detaching from the substrate. Strikingly, detaching from the substrate results in a local collapse of the periodic myofibrils (Figure 5F and G, Figure 5 – video 2). This further supports our hypothesis that mechanical tension is the long-range signal that instructs and coordinates sarcomere assembly across the entire myofiber.

### Mechanical tension build-up precedes sarcomere assembly

In order to assess the role of tension in triggering the transition from a dynamic cytoskeleton in extending myotubes to periodic myofibrils in stable myofibers, we quantified mechanical tension using laser nano-surgery. We severed entire myotubes and myofibers expressing either endogenously tagged α-actinin2 (Figure 6A, myotube and immature myofiber), or titin (Figure 6A, transitional and mature myofibers). The recoil can be visualised by plotting the position of the severed ends as kymographs over time (Figure 6A, Figure 6 – video 1). After cutting, severed myotubes, as well as immature, transitional and mature myofibers all displayed a recoil response of their severed ends. We conclude that mechanical tension is present along the long axis of myotubes and myofibers of all morphological categories, thus preceding sarcomere assembly and also persisting during sarcomere maturation.

**Figure 6.**
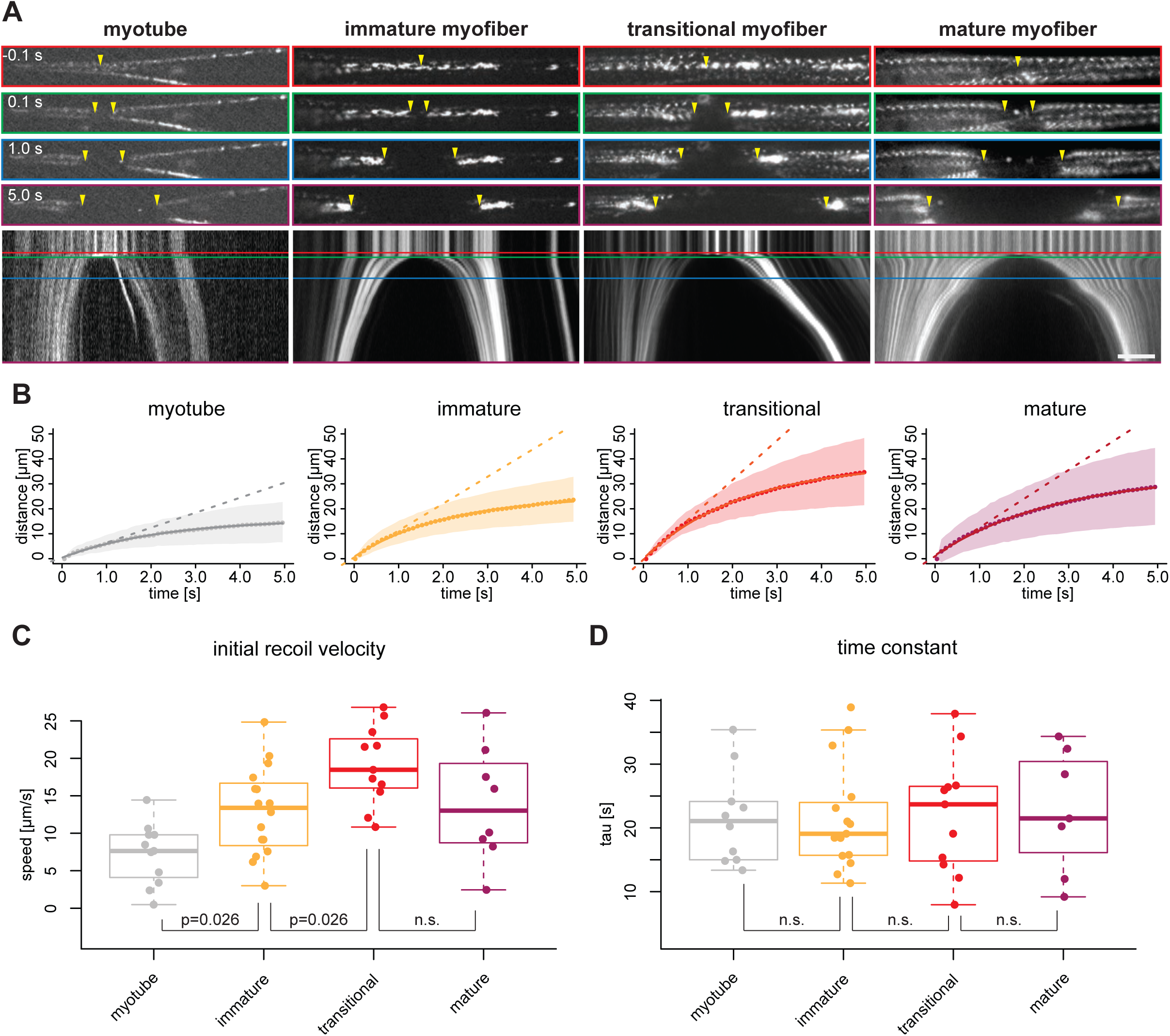
Mechanical tension precedes and persists during sarcomere formation. **A)** Line laser ablation experiments for each morphological category (myotubes, immature, transitional and mature myofibers). Myotube and immature myofibers were labelled with mKate2-α-actinin2, while transitional and mature myofibers were labelled with GFP-titin. Top, upper middle, middle, and lower middle panels show 0.1 s prior to, 0.1 s, 1.0 s, and 5 s post ablation, respectively. Bottom panel shows a kymograph of ablated myofibers. Coloured boxes correspond to coloured lines in the kymograph. Yellow arrowheads show sites of ablation and severed ends. **B)** Quantification and mathematical modelling of recoil distances. Dots and shaded area show average and standard deviation of recoil distance, respectively. Dashed straight lines show a linear model fitted to estimate the initial recoil velocity. Solid curve shows viscoelastic model fitted to estimate the relaxation time τ. **C-D)** Box plot of initial recoil velocity (C) and relaxation time (D) from ablation experiments in each morphological category: n = 11 (myotube), 16 (immature), 11 (transitional) and 8 (mature). Black bars mark the distance between 1^st^ and 3^rd^ quadrant. White dots mark the median values. Multiple pair-wise t-test: n.s.: p>0.05. Scale bar: 10 µm. **[Related to Figure 6 – figure supplement 1, figure supplement 2, Figure 6 – video 1, video 2 and video 3. Source data available for Figure 6C and D.]**

Additionally, we found that transitional and mature myofibers frequently show fast motions as consequences of active contractions (Figure 6 – figure supplement 1A, B and Figure 6 – video 2). Similar active contractions have been observed when severing immature myofibrils in *Drosophila*, as a consequence of laser-induced calcium release from the sarcoplasmic reticulum (Weitkunat et al., 2017) and have also been observed spontaneously in mature human myofibers after applying the same differentiation protocol (Tanoury et al., 2021). This demonstrates that transitional myofibers harbour already contractile sarcomeres.

If mechanical tension build-up is a trigger for periodic myofibril assembly, one would expect higher tension levels in transitional myofibers compared to earlier stages. As it is well established that the recoil velocity speed after the cut correlates with the amount of mechanical tension (Behrndt et al., 2012; Hutson et al., 2003), we tracked the recoiling ends of severed myotubes and myofibers (excluding the actively contracting ones) and fitted both a linear model and a viscoelastic model (Figure 6B) to calculate the initial recoil velocity (Figure 6C) and the viscoelastic relaxation time τ (Figure 6D), respectively. τ is comparable between all morphological categories, suggesting that they have comparable material properties. However, immature and transitional myofibers show a progressively higher initial velocity compared to myotubes (Figure 6C), demonstrating that they are under higher tension levels.

To directly probe tension within the assembling myofibrils, we performed nano-lesions to severe only individual myofibrils without affecting the rest of the cell, taking advantage of the high spatial precision of the pulsed infrared laser (see methods). We found that severed myofibrils are recoiling, demonstrating that they indeed are under tension (Figure 6 – figure supplement 2, Figure 6 – video 3). This strongly suggests that the assembling and maturing myofibrils are the major source of the mechanical tension present in the myofibers. We found that severing a mature myofibril leads to the collapse of its neighbouring sarcomeres (Figure 6 – figure supplement 2, Figure 6 – video 3), similar as in detaching myofibers observed in Figure 5F. In conclusion, these data demonstrate that attached myofibers build up mechanical tension prior to and during sarcomere formation, supporting our hypothesis that tension coordinates the assembly of long periodic myofibrils upon surpassing a threshold. These myofibrils need stable anchoring to prevent them from collapsing.

### Force-resistant attachment results in myofiber end clustering

*In vivo*, myofibers are stably attached to tendons cells that connect to the skeleton (Lemke and Schnorrer, 2017; Schweitzer et al., 2010). To mechanically anchor the myofibrils, the terminal Z-disc of each myofibril is connected to the cytoplasmic tails of integrins by a specialised actin-rich zone (Green et al., 2018; Lemke et al., 2019; Sun et al., 2019). Integrins in turn are localised to the cell membrane and their extracellular domains bind to various components of the muscle-tendon extracellular matrix (ECM) (Brown, 2000; Hynes, 2002). As our cultures do not contain prepatterned tendon-based attachments, we wondered how myofiber bundles develop stable attachments around day 7?

Integrins function as *α*ß-dimers. The human genome contains 18 *α*- and 8 ß-integrin subunits, forming 24 functionally distinct dimers (Ringer et al., 2017; Takada et al., 2007). To identify which ones are likely relevant for attachment of muscle fibers in our cultures, we made use of the developmental gene expression time-course and found that the fibronectin binding *α*vß1-, *α*5ß1- and *α*6ß1-integrins are the highest expressed integrins throughout the time course (Supplementary Table 1). Additionally, *α*7-integrin is strongly upregulated at day 7, forming with ß1 a laminin receptor (*α*7ß1-integrin) (Hynes, 2002; Plow et al., 2000). These expression data fit with the severe muscle fiber and sarcomere assembly phenotypes of ß1-integrin knock-out mice (Brakebusch and Fassler, 2005; Schwander et al., 2003) and the muscular dystrophy phenotype of *α*7-integrin knock-out mice (Mayer et al., 1997). Hence, we used ß1-integrin as main adhesion marker together with paxillin, which is a force-sensitive component of adhesions (Theodosiou et al., 2016) and co-stained our cultures for titin and actin (Figure 7A-D and Figure 7 – figure supplement 1A-D). Consistent with our transcriptomics data, we found that ß1-integrin protein levels increase when the myofibers differentiate, coinciding with an increase in titin and actin levels (Figure 7E-G).

**Figure 7.**
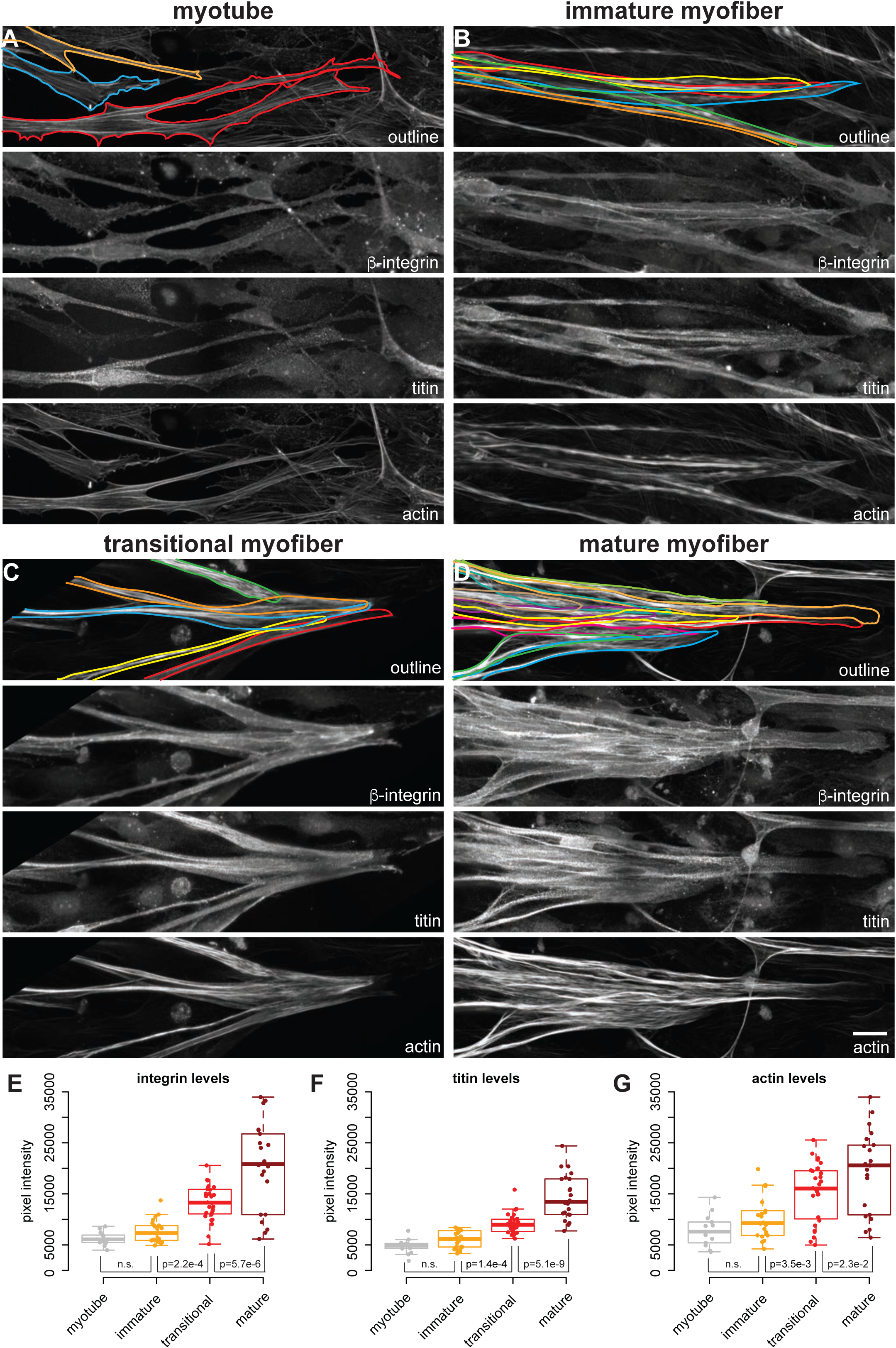
Attachment morphology during myofiber bundle maturation. **A-D)** Immunostainings of myofibers with β1-integrin (upper middle), titin (lower middle) and actin (stained with phalloidin, bottom) in myotubes (A), immature (B), transitional (C) and mature myofibers (D). The top panel shows the outline of the myofibers superimposed on the actin channel. E-G) Intensity of β1-integrin (E), titin (F) and actin (G) signals measured with a 50-pixel-wide line-ROI from myotubes (n = 13), immature (n = 22), transitional (n = 28) and mature myofibers (n = 22). Scale bars: 20 µm. **[Related to Figure 7 – figure supplement 1. Source data available for Figure E-G.]**

Closer inspection of high-resolution images showed that ends of myotubes contain an actin-rich network forming prominent filopodia with some integrin and paxillin-based focal adhesions (Figure 7 – figure supplement 1A). This is consistent with the highly dynamic extension behaviour of myotube ends that we documented by live imaging (Figure 1 – figure supplement 1B), resembling the dynamic focal adhesions of migratory cells (Wehrle-Haller, 2012). Similar integrin and paxillin foci are detected at the ends of immature myofibers, however integrin also localises along the entire outline of the immature myofiber (Figure 7B, Figure 7 – figure supplement 1B). This suggests that the immature myofibers probe their environment over large areas, which is consistent with their above describe dynamics.

When the myofiber bundle foci stabilise and myofibers convert to transitional myofibers, we found a very notable concentration of integrin and paxillin at the interface of neighbouring myofibers (Figure 7C, Figure 7 – figure supplement 1C). This fiber-fiber localisation of integrin and paxillin further increases in mature myofibers, which frequently form large clustered foci (Figure 7D, Figure 7 – figure supplement 1D). To better document the fiber-fiber proximity we visualised the myofiber outlines with the cell surface marker NCAM1 and confirmed that immature as well as mature myofibers form immediate contacts at the bundle foci (Figure 7 – figure supplement 2).

Taken together, our data suggest that the myofiber bundle foci of transitional and mature myofibers are indeed stabilised by force-resistant integrin-based fiber-fiber attachments to the surrounding extracellular matrix. These stable attachment foci can counteract the high tension produced in each transitional myofiber driving sarcomere assembly, and thus sarcomere assembly is coordinated with myofiber bundling at the supra-cellular level.

## Discussion

### Multi-scale self-organisation in human myofibers *in vitro*

*In vivo,* human muscle tissue displays a very ordered organisation ranging from the molecular scale of sarcomeric proteins to the tissue scale of myofiber bundles. This ensures that molecular forces produced by myosin motors translate efficiently into body movement. Here, we used a 2D-culture system to investigate how muscle cells can organise themselves in the absence of any external cues. We discovered that myofibers show an autonomous multi-scale self-organisation when cultured *in vitro* (Figure 8).

**Figure 8.**
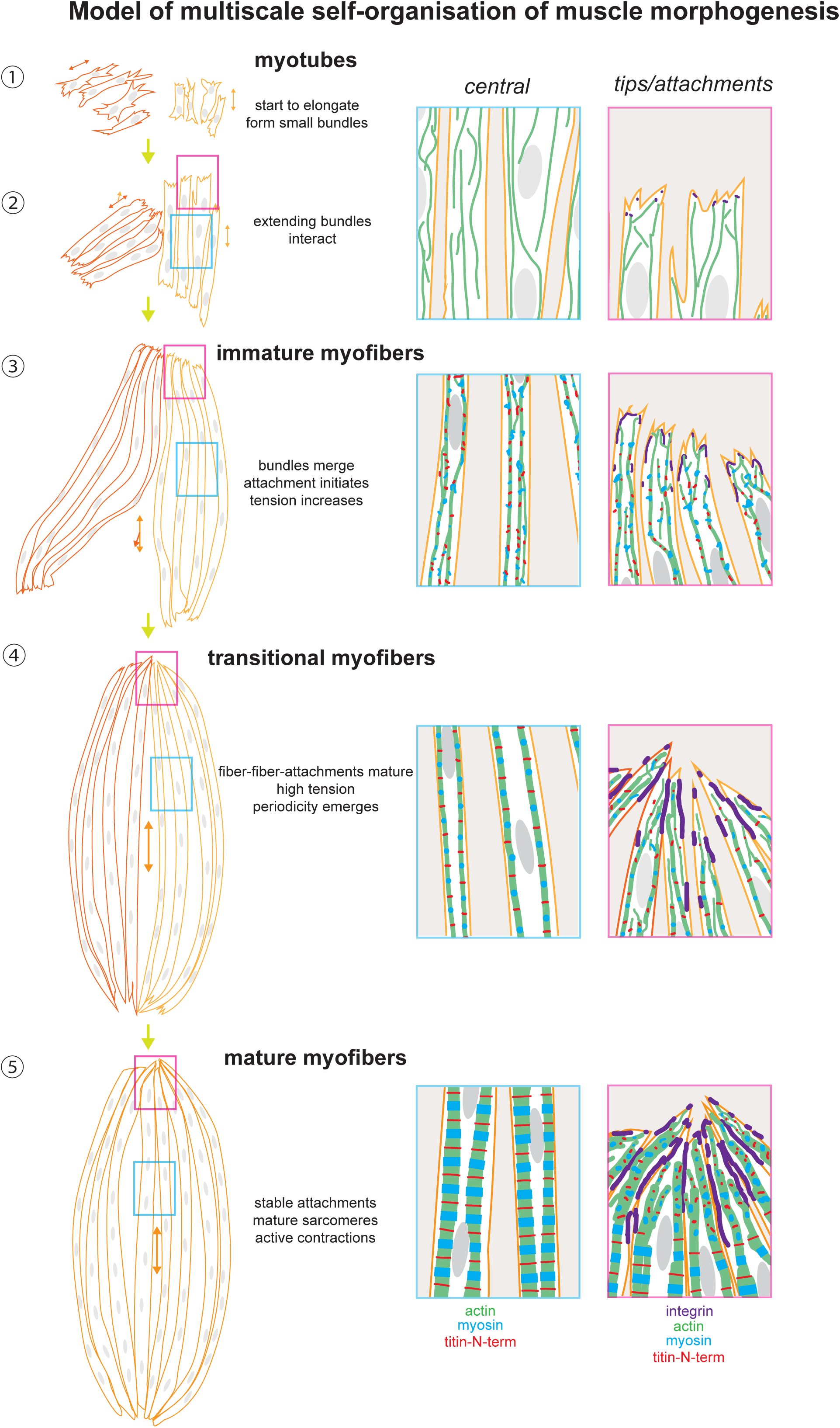
A tension driven self-organisation model of human skeletal muscle morphogenesis across tissue, cell and molecular scales. See text for description.

At the tissue scale, myofibers self-organise into large bundles sharing one long axis, the future muscle contraction axis. Most myofibers within one bundle stably connect to one shared attachment focus on either end. This resembles the *in vivo* situation of muscle fiber bundles growing in the same direction and connecting to shared tendon and skeletal elements (Schweitzer et al., 2010; Zelzer et al., 2014).

At the cellular level, myotubes first elongate actively, then stabilise to a defined length, likely depending on the culture and matrix conditions (Jensen et al., 2020). At this stage, myofiber ends start to establish stable connections with their environment, coinciding with their transition from immature to transitional myofibers. Again, this behaviour recapitulates the *in vivo* situation, in which myotubes first elongate towards their future tendons, then they stably attach to them, before they organise their contractile myofibrils (Gros et al., 2004; Kardon et al., 2003; Schnorrer and Dickson, 2004; Weitkunat et al., 2014).

At the molecular level, sarcomeres emerge homogeneously with first periodic regions distributed across a long part of the myofiber after its stable attachment. These immature myofibrils then mature into chains of hundred or more periodic sarcomeres that span most of the mature myofiber. Further maturation will result in the lateral expansion of the periodic myofibrils by lateral alignment of neighbouring myofibrils and incorporation of newly recruited components. A similar simultaneous myofibril assembly and maturation mechanism has been described in *Drosophila* muscles *in vivo* (Weitkunat et al., 2014; 2017). Hence, our human muscle *in vitro* system does recapitulate various aspects of muscle morphogenesis *in vivo*.

The alignment of neighbouring myotubes to a shared contraction axis has been also found when cultured C2C12 mouse myoblasts were differentiated on patterned substrates with a defined axis (Guillamat et al., 2020; Junkin et al., 2011). The alignment of myofibers in this system depends on the compliance of the substrate, with very soft hydrogels resulting in a highly parallel myotubes that do not converge to shared attachment foci (Jensen et al., 2020). Hence, self-organised alignment appears to be a general phenomenon of differentiating myotubes. However, prior to our work it was unknown how myotube alignment at the tissue scale coordinates with muscle and sarcomere morphogenesis at cell and molecular scales in human muscle.

Here we have presented evidence strongly supporting mechanical tension as one key signal to coordinate the self-organisation across scales. First, we detected different levels of mechanical tension during the time-course of myofiber differentiation, starting with low tension in myotubes and resulting in high tension in the assembled transitional and mature myofibers. Second, we demonstrated that this tension is present across the assembling myofibrils. Since myofibrils span the entire muscle fiber, tension likely communicates rapidly across scales. Third, we found a strong correlation of tension increase with maturation of integrin attachments at the myofiber bundle foci. It is well established in many cell types that mechanical load on integrin adhesions exerted by the actin cytoskeleton results in their maturation to better withstand increased forces (Austen et al., 2015; Bulgakova et al., 2017; Lemke et al., 2019; Sun et al., 2019). Finally, we found that releasing tension either acutely by laser cutting a myofibril or by long-term detachment of a myofiber tip results in a collapse of the periodic myofibrils, which will likely result in a disassembly of the ordered structure. Altogether, this strongly suggests that mechanical tension is a key signal that coordinates morphogenesis in myofibrils, myofibers and myofiber bundles.

### A self-organisation model across scales is coordinated by tension

How may tension coordinate muscle morphogenesis across scales? In our working model (Figure 8), myotubes elongate during the initial culture phase via protrusive ends. At this stage, they are already forming first mechanical contacts with neighbouring myotubes using filopodial extensions, which in response align to form a small and short myotube cluster sharing a common axis. Tension level is low and sarcomeric genes are not or little expressed (Figure 8 – steps ①, ②). As myotubes elongate further, they convert to immature myofibers: sarcomeric proteins become expressed and actin filaments assemble into continuous myofibrils that are under higher tension but are still not periodic. The elongating myofibers start to merge with neighbouring bundles to form a common axis and shared attachment foci (Figure 8 – step ③). The stable attachment enables a further increase in tension and the now high levels of sarcomeric proteins can assemble into periodic myofibrils throughout most of the myofiber (Figure 8 – step ④). The self-organisation process is likely myosin-driven, with myosin being responsible for the tension build-up and the ordering of the sarcomeric components, as has been shown in *Drosophila* (Dasbiswas et al., 2018; Loison et al., 2018) as well as in cardiomyocytes *in vitro* (Chopra et al., 2018; Fenix et al., 2018). Hence, we propose that tension driven myofibril self-organisation is evolutionarily conserved across muscle types and tension can act as a compass to orient and coordinate between all the molecular components for coordinated assembly into continuous periodic myofibrils (Lemke and Schnorrer, 2017).

Importantly, we found that integrin-based attachments are concentrated at the ends of the myofibers, in particular at the contact interfaces between neighbouring myofibers. Likely these integrins are mechanically engaged as is the case at muscle attachments *in vivo* (Charvet et al., 2012) thus allow the bundling of the myofibers to large bundles with one large attachment focus at each end. Such, each of the foci couples mechanically to the extracellular matrix to resist the active myosin-based contractions that become prominent in mature myofibers (Figure 8 – step ⑤). Such active contractions are also essential for myofibril maturation in *Drosophila* (Weitkunat et al., 2017), primary chick myofibers cultured *in vitro* (Kagawa et al., 2006) and in turn even for tendon and bone maturation *in vivo* (Felsenthal and Zelzer, 2017; Shwartz et al., 2012). Taken together, our model proposes a key role for mechanical tension to coordinate the self-organisation of muscle morphogenesis across scales.

### Tension sensing

Which molecules may sense tension to signal a coordinated assembly of myofibrils and myofibers into bundles? It is likely that tension is sensed at the myofiber level, as sarcomeres appear homogenously across the entire lengths of myofibers. A myosin-dependent self-organisation of ordered periodic actomyosin structures has even been observed in non-muscle cells (Dasbiswas et al., 2018; Hu et al., 2017). However, in contrast to non-muscle cells, the sarcomeric striations are stable over long time periods and have period length of 2 to 3 micro meters. This is achieved by titin, which ‘locks’ actin and myosin filaments in place, fulfilling its established ruler function (Brynnel et al., 2018; Granzier et al., 2014; Linke, 2018; Tskhovrebova and Trinick, 2003). In turn, titin is the major source of passive tension present in mature muscle and is responsible for the mechanical integrity of sarcomeres (Li et al., 2020; Rivas-Pardo et al., 2020; Swist et al., 2020). Hence, it is conceivable that the giant elastic titin molecules may serve as tension-sensitive catch bonds, whose incorporation into the myofibril subsequently triggers spatial segregation of actin and myosin domains into regular sarcomeres. However, molecular forces across titin during myofibril assembly have not yet been measured directly.

Another possible mechanism may exist at the ends of myofibers, where integrin-based attachment sites are mechanically linked to the actin cytoskeleton. Since myofiber elongation is mainly driven by actin dynamics of the myofiber ends (Schnorrer and Dickson, 2004; Weitkunat et al., 2014), a tension-sensing mechanism located at the myofiber ends could transmit the tension signal via the actin cytoskeleton to negatively regulate dynamics of myofiber ends and initiate stable attachment (Richier et al., 2018). Recent work from *Drosophila* muscle attachments revealed that high mechanical load on integrin does indeed recruit large amounts of the integrin adaptor talin, bridging from integrin to the actin cytoskeleton and the first sarcomeric Z-disc (Bulgakova et al., 2017; Green et al., 2018; Luis and Schnorrer, 2021; Moser et al., 2009). The large amounts of talin can then share the high mechanical load amongst all the talin molecules to achieve stable long-term attachment (Lemke et al., 2019). While we have not been able to detect costameric localisation of integrins in our cultured myofibers, likely due to their lack of maturity, we do not exclude a role of costameric integrins in mediating mechanical forces and promoting sarcomere formation (Schwander et al., 2003; Sparrow and Schöck, 2009), since it has been shown that costameres in mature myofibers play important roles in transmitting forces from the sarcomere to the sarcolemma and the extracellular matrix (Peter et al., 2011).

Finally, it is important that the transcriptional status of the differentiating myofiber fits with its morphological differentiation. Thus, tension may also feedback on the transcriptional machinery to boost expression of the sarcomeric components that are needed in large amounts to assemble myofibrils but are not present in extending myotubes. Such a feedback mechanism has recently been proposed in developing flight muscles, which use the mechanosensitive Hippo/YAP pathway to boost sarcomere gene expression before and during myofibril morphogenesis (Kaya-Copur et al., 2021). Tension could also be sensed directly at the nuclear envelope, which is linked by the LINC complex to the actin cytoskeleton, inducing changes of nuclear shape and position during myofiber differentiation (Janota et al., 2020; Wang et al., 2018). Hence, it is conceivable that in the developing human muscles tension is sensed to boost sarcomeric gene expression. Such a mechanism would ensure that sarcomeres only assemble from their components when the myofibers are properly attached.

### Making even better muscle *in vitro* – towards a muscle-skeleton ‘organoid’?

The 2D myofibers generated *in vitro* are still immature compared to large adult human muscle fiber bundles attached to tendons and bone *in vivo*. Our myofibers show highest expression of embryonic (MYH3) and neonatal (MYH8) myosin isoforms (see Supplementary Table 1) (Schiaffino et al., 2015; Tanoury et al., 2021) and their myofibrils remain rather thin. However, longer-term culture often results in myofiber detachment followed by degeneration, likely because the attachment sites are not stable enough to withstand the even higher forces produced by more mature fibers. This could be partially solved by overlaying with second layer of Matrigel to provide additional mechanical support in 3D (Roman et al., 2018). The successful differentiation of contractile human neuromuscular organoids from neuromesodermal progenitors in 3D (Faustino Martins et al., 2020) also suggests that three-dimensional support of the myogenic cultures is advantageous.

In summary, our current study has uncovered a remarkable transition from an initial population of dynamic, elongating myotubes to stable and contractile bundles of myofibers that is likely coordinated across tissue, cell and molecular scales via mechanical tension. Given that muscle-generated mechanical forces are crucial for optimal patterning and morphogenesis in bones and tendons (Felsenthal and Zelzer, 2017), it is tempting to hypothesise that mechanical tension may also coordinate the morphogenesis across muscle and connective tissue lineages, resulting in a self-organised musculoskeletal system. Introducing progenitors of tendon or other connective tissues at defined positions of the cultures will be a next step towards a minimal sufficient system that can recapitulate mature muscle-tendon-skeleton development *in vitro*.

## Materials and Methods

### Human iPSC-derived 2D skeletal myoculture

The following human iPSC lines were used to perform myogenic differentiation: untagged NCRM-1 (RUCDR, Rutgers University), GFP-titin (derived from PGP-1, (Sharma et al., 2018)) and mKate2-α-actinin2 (derived from NCRM-1, this study, see below).

Maintenance of human iPSCs and their primary myogenic differentiation were performed as described in (Chal et al., 2016; Tanoury et al., 2021). For secondary differentiation, primary myogenic cultures derived from iPSCs were dissociated at 3 weeks-post-differentiation, filtered through a 70-μm and a 40-µm cell strainer, and cryopreserved in custom cryomedia (Chal et al., 2016). After thawing in 37°C water bath, myogenic progenitors were resuspended in skeletal muscle growth medium (SKGM-2, Lonza, Cat. CC-3245) with 10 µM ROCK inhibitor, and seeded at a density of 5-10 k/cm^2^ onto polymer coverslip-bottomed imaging dishes (culture-insert 4 well in µ-dish 35 mm, ibidi®, Cat. 80466 for fixed experiments or micro-insert 4 well in µ-dish 35 mm, ibidi®, cat. 80406 for live experiments) coated with Matrigel (Corning, Cat. 354277). After 24 h, culture medium was replaced by SKGM-2 medium without ROCK inhibitor. Myogenic progenitors were allowed to proliferate for 1-2 days until reaching ∼80% confluence, then induced for myogenic differentiation by replacing SKGM-2 medium with the KCTiP differentiation medium (Tanoury et al., 2021): DMEM/F12 medium (Thermo Fisher Scientific, Cat. 11320033) supplemented with 2% knock-out serum replacement (Invitrogen, Cat. 10828028), 1% ITS (Life Technologies, 41400045), 0.2% Pen/Strep (Life Technologies, Cat. 15140122), 1 µM Chiron (Tocris Bioscience, Cat. 4423), 10 μM TGF-β inhibitor SB431542 (Tocris, Cat. 1614), and 10 μM of prednisolone (Sigma-Aldrich, Cat. P6004). KCTiP differentiation medium was subsequently changed on days 1 and 2, and then was refreshed every other day until the end of the cell culture.

### Generation of the mKate2-α-actinin2 knock-in reporter line

Two different sgRNA targets were designed using the CRISPOR tool (http://crispor.tefor.net) surrounding the start site of human ACTN2 gene for N-terminal tagging (primers sequences are listed below). sgRNAs oligonucleotides were annealed and cloned into pSpCas9(BB)-2A-GFP. Donor plasmid was designed by NEBuilder (https://nebuilderv1.neb.com) using Gibson assembly in pUC10 backbone. 0.8-1 kb nucleotide sequence before the sgRNA cutting site as 5’-homology arm (HA), mKate2 and 0.8-1 kb nucleotide sequence after the sgRNA cutting site as 3’-homology arm were used to generate primers for cloning (see below). A silent mutation was introduced on each of the donor template (5’-HA and 3’-HA) in the PAM site to prevent the CRISPR from targeting the recombined sequences. 5’-HA and 3’-HA were PCR amplified using human genomic DNA. mKate2 dsDNA was synthesized and obtained from Integrated DNA Technologies. Gibson cloning method was used to ligate each part to construct the full donor plasmid (New England Biolabs).

#### Primer sequences

*sgRNA1*

sgRNA target site: CACGTAGTTGTACTGCACGC

Forward Primer: 5’CACCGCACGTAGTTGTACTGCACGC3’

Reverse Primer: 5’AAACGCGTGCAGTACAACTACGTGC3’

*sgRNA2*

sgRNA target site: ACTGCACGCCGGGCTCTATC

Forward Primer: 5’CACCGACTGCACGCCGGGCTCTATC3’

Reverse Primer: 5’AAACGATAGAGCCCGGCGTGCAGTC3’

*5’HA (R)* (anneals pUC19): GACGAAAGGGCCTCGTGATAC

pUC19 (F)(anneals 5’HA): TATCACGAGGCCCTTTCGTCGAGACACGAGAGAGAGAG

mKate2 (R) (anneals 5’HA): CGCTCACCATCATGTACTCATCCTCGTC

5’HA (F) (anneals mKate2): TGAGTACATGATGGTGAGCGAGCTGATTAAG

3’HA (R) (anneals mKate2): ACTCCTCCTCCTGGATTCATCTGTGCCCCAGTTTG

mKate2 (F) (anneals 3’HA): GCACAGATGAATCCAGGAGGAGGAGTGG

pUC19 (R) (anneals 3’HA): GTCATCACCGAAACGCGCGAAAAAAGTTCAGAGCGTTTGG

3’HA (F) (anneals pUC19): TCGCGCGTTTCGGTGATG

### Immunostaining of fixed myocultures

Myocultures in imaging dishes were rinsed one time in 1× PBS before fixed in 4% PFA for 20 minutes at room temperature. Fixed myocultures were rinse one more time in 1× PBS, permeabilized in 1× PBS supplemented by 0.1% Triton X-100 (Sigma-Aldrich, Cat. T8787) for 15 minutes at room temperature, then incubated in blocking buffer composed of 1×PBS supplemented with 10% Gibco Fetal Bovine Serum (Thermo Fisher Scientific, Cat. A4766801) and 0.1% Triton X-100. Myocultures were subsequently incubated with appropriate primary antibodies diluted in blocking buffer at appropriate concentrations overnight at 4°C. Myocultures were then washed three times with 1× PBS supplemented by 0.1% Tween20, and incubated with appropriate fluorescently labelled secondary antibodies and Phalloidin diluted 1× PBS supplemented by 0.1% Tween20 for 2 hours at room temperature. Myocultures were then washed three times with 1× PBS supplemented by 0.1% Tween20 (Sigma-Aldrich, Cat. P9416). The culture inserts were removed to allow mounting with a top coverslip and appropriate mounting media. Samples were sealed with nail polish and stored at 4°C until imaging.

The following primary antibodies were used: rabbit-anti-titin (N-term) (Sigma, Cat. HPA007042, 1:50), mouse anti-MyHC (DSHB, Clone A4.1025, 5 ng/µL), rat anti-β1integrin (DSHB, Clone AIIB2, 5 ng/µL), mouse anti-paxillin (Invitrogen, Cat. AHO0492, 1:5), mouse-anti-NCAM1 (DSHB, Clone 5-1H11, 5 ng/µL). The following secondary antibodies were used: goat anti-mouse Alexa405 (Invitrogen, Cat. A31553, 1:250), goat anti-rabbit Alexa405 (Invitrogen, Cat. A31556, 1:250), goat anti-rabbit Alexa488 (Life Technologies, Cat. A11034, 1:250), goat anti-rat Alexa488 (Invitrogen, Cat. A11006, 1:250), goat anti-mouse Alexa488 (Invitrogen, Cat. A 11001, 1:250), goat anti-mouse Alexa633 (Life Technologies, Cat. A21052, 1:250), goat anti-rat Alexa633 (Life Technologies, Cat. A21094, 1:250), goat anti-rabbit Alexa633 (Invitrogen, Cat. A 21070, 1:250). The following staining reagents were used: rhodamine-phalloidin (Invitrogen, Cat. R415, 1:500), Alexa568-phalloidin (Invitrogen, Cat. A12380, 1:500). The following mounting media were used: Prolong Gold Antifade without DAPI (Invitrogen, Cat. P36934), Prolong Gold Antifade with DAPI (Invitrogen, Cat. P36935).

### Fluorescence imaging of fixed myocultures

Myocultures were differentiated in polymer coverslip-bottomed imaging dishes (culture-insert 4 well in µ-dish 35 mm, ibidi®, cat. 80466) coated with Matrigel and fixed as described above. Fluorescent images of fixed myocultures were acquired on a Zeiss LSM880-I-NLO inverted confocal microscope using the following objectives: EC Plan-Neofluar 10x/0.3, Plan-Apochromat 100x/1.4 Oil DIC. For images acquired at 10x magnification, 6×6 tiles at zoom 1.0 were used to cover the entire culture well, while single z-slices were acquired. For images acquired at 100x magnification, various tiles at zoom 1.0 were used to cover a larger area, with a z-step size of 0.3-0.5 µm.

### Phase-contrast time-lapse imaging of live myocultures

Myocultures were differentiated in polymer coverslip-bottomed imaging dishes (micro-insert 4 well in µ-dish 35 mm, ibidi®, cat. 80406) coated with Matrigel as described above. Phase-contrast time-lapse images of live myocultures were acquired on a Zeiss Axio Observer.Z1 inverted wide-field microscope with 5% CO_2_ at 37°C using an EC Plan-Neofluar 20x/0.5 objective. 4×4 tiles at zoom 1.0 were used to cover a larger area. Images were acquired every 30 minutes, for up to 6 days. Closed culture dishes filled with ample media kept in a moisturized imaging chamber allowed for stable long-term imaging without perturbing the sample by media change. Software autofocus was used to keep the imaging plane constant.

### Fluorescence time-lapse imaging of myofibrillogenesis

Myocultures were differentiated in polymer coverslip-bottomed imaging dishes (micro-insert 4 well in µ-dish 35 mm, ibidi®, cat. 80406) coated with Matrigel as described above. Fluorescent time-lapse images of live myocultures were acquired on an Olympus IX83 inverted microscope equipped with a Yokogawa CSU-Wi spinning disc scanner unit using an UPlanXApo 60x./1.42 oil immersion objective. An Okolab UNO-T-H-CO2 heated stage was used to carry on incubation with 5% CO_2_ at 37°C during the entirety of time-lapse experiment. Tiled z-stacks were acquired every 15 minutes with a range of 10 µm at a step size of 1 µm, for up to 3 days. Closed culture dishes filled with ample media allowed for stable long-term imaging without perturbing the sample by media change. Z-drift compensation was used to keep a reference plane constant.

### Laser-ablation of live myofibers

Laser-ablation and fluorescent live imaging of live myocultures were performed on a Nikon Eclipse TE 2000-E inverted spinning disc microscope (Ultraview ERS, Perkin Elmer) and a near-infrared (NIR, 1030 nm) femtosecond laser at 50 MHz repetition rate (t-Pulse, Amplitude Systems) (Collinet et al., 2015). Whole myofiber cuts were achieved by focusing the NIR laser using a 100×/1.4 oil-immersion objective with an average power of 500 mW at the back aperture, then moving the focused beam along a 5∼10 µm line perpendicular to the target myofiber at a speed of 300 µm/s using an xy-galvanometer (Cambridge Technologies). Fluorescent time-lapse images were acquired every 100 milliseconds prior to, during and post-ablation for up to 10 seconds.

### Transcriptomics analysis

The bulk RNA-seq data analysis was performed on the already analysed data available under the GEO accession number GSE164874. Starting from the raw count matrix file “GSE164874_SecondaryDifferentiation_raw_counts.xlsx”, we kept here only the relevant conditions, namely the three replicates for the SKGM-2, KCTiP at 7 days and KCTiP at 15 days of differentiation. In order to compare each time point to a baseline level we artificially generated three pseudo-replicates for the baseline condition. For each of the three pseudo-replicates we computed a baseline value by averaging different replicates of the three time points. Raw count matrix filtered for the conditions of interest and with pseudo-replicates can be found in the Supplementary Table 1. Normalization, rlog transformation and differential expression analysis were done in the R environment (v3.6.0), using the DESeq2 package (v 1.22.2) (Love et al., 2014). The output files of the DESeq2 analysis can be found in the Supplementary Table 1.

Cluster analysis was performed on the normalized counts using the Mfuzz R package (v 2.42) (Kumar and Futschik, 2007). The eliminate the noise, genes with lower than 50 total reads were filtered out. The data have been partitioned into 10 clusters according to the Dmin function provided by Mfuzz. Genes with membership values lower than 0.2 were filtered out generating non overlapping clusters. Moreover, we performed 10 clustering runs and evaluated the stability of the clusters by computing the Jaccard Index. Cluster profiles are displayed in Figure 3A, while the gene membership lists in the clusters are available in Supplementary Table 1. We performed GO enrichment of each cluster making use of Enrichr (Kuleshov et al., 2016). Results of the enrichment analysis can be found in the “GO_enrichment_clusters.xlsx” supplementary file.

### Image analysis and quantification

#### Segmenting and quantifying myofibers and nuclei

Myofibers were manually segmented from 10× phalloidin channels as freehand line ROIs in Fiji (Schindelin et al., 2012) by tracing with a stylus on a digital drawing pad. Length of these line ROIs were then measured in Fiji. Width of myofibers were manually quantified from 100× phalloidin channels as length of line segment ROIs perpendicular to the myofibers in Fiji.

To count nuclei automatically, freehand line myofiber ROIs generated from 10× phalloidin channels were reapplied to the DAPI channel to quantify pixel intensities along the line ROIs. A custom R-script was developed to extract the numbers and positions of local maxima in DAPI pixel intensities above arbitrary thresholds as nuclei signal. The results of automatic counting were compared with manual counting for selected images and the two were similar.

Quantifications of β1-integrin, titin, and actin levels for Figure 7 were performed using line ROIs of a width of 50 pixels (4 µm) on multi-channel maximum projection images with a depth of 5 µm. Average pixel intensity per line ROI was then taken as a measure of relative protein levels.

#### Segmenting and quantifying myofiber bundles and attachment foci

First, myofibers were manually segmented from 10× phalloidin channels as freehand line ROIs in Fiji by tracing with a stylus on a digital drawing pad. Then, line ROIs were grouped as bundles with a custom script in R using a simple clustering algorithm based on the Hausdorff distance (Borchers, 2021). The Hausdorff distance is defined as the maximum distance of a set of points to their nearest point in the other set (Rote). The Hausdorff distance is chosen to quantify the distance between two line-ROIs, as it takes into account every point and is thus superior than simply comparing average distances of line ROIs. Briefly, each set of line ROIs is represented by a set of x,y coordinates. A Hausdorff distance is then calculated between each sets of x,y coordinates to obtain a distance matrix between pairwise line ROIs. Line ROI pairs with Hausdorff distances below arbitrary thresholds are grouped into one myofiber cluster. One unique distance threshold is chosen for each time point, based on best matching between automatically generated clusters with visual inspections. Once segmented, the number of myofiber clusters is automatically counted. The area of each myofiber cluster is automatically quantified as the convex hull of all x,y coordinates in line ROIs within that cluster.

Attachment foci are manually segmented from 10× phalloidin channels, as sets of x,y coordinates where myofiber bundle converges.

#### Vector field analysis of myofiber bundles

Vector field analysis of myofiber bundles was performed using the OrientationJ plugin (Rezakhaniha et al., 2012) in Fiji (Schindelin et al., 2012) combined with custom R-scripts. First, in order to remove edges and empty space without myoculture, a central window of 3200×3200 pixels was cropped from the original 5000×5000 pixels images from 10× phalloidin channels. Then, the cropped image was analysed using the OrientationJ vector field plugin with the following parameters: structure tensor local window σ = 10 pixels, vector field grid size = 100 pixels. The output from OrientationJ (Figure 2 – figure supplement 1A-F) is further analysed with custom scripts in R by adapting algorithms from (Guillamat et al., 2020) as follows:

The spatial correlation *C* (Figure 2 – figure supplement 1G) was defined as

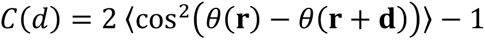

where *θ* (**r**) is the local orientation angle of the vector field at a grid position **r**, **d** is the separation vector to a second grid position **r**+**d** at distance |**d**|=*d*, and brackets 〈 〉 denote an average over all pairs of grid points with same distance *d*. The characteristic nematic length η_L_ was defined as the intersection of the initial linear decay of *C*(*d*) and the *x* axis (Figure 2 – figure supplement 1G).

The local nematic order parameter *S* (Figure 2 – figure supplement 1A’-F’) was defined as

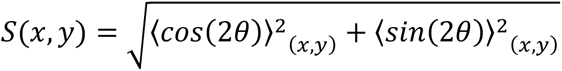

where 2 is again the local orientation angle of the vector field at a given grid position, and brackets 〈 〉 denote an average over a local regions of interest with center position (*x*,*y*). Specifically, we used square regions of interest comprising 4x4 grid positions each. The local nematic order parameter *S* was plotted as a heatmap (Figure 2 – figure supplement 1A’-F’). The fraction of disorder parameter is calculated as the fraction of local regions of interest for which the local nematic order parameter *S* is below a threshold of 0.5 (Figure 2 – figure supplement 1I).

#### Segmenting myofibrils and generating intensity profiles

1D ROIs of myofibrils were manually generated from 3D stacks of 100× titin channel in four steps (Figure 4D): First, a spline is drawn on the xy plane of the original 3D stack. This xy-spline is used to generate a xy-straightened 3D volume (①). Second, the xy-straightened 3D volume is projected onto the xz plane, where a second spline is drawn. This xz spline is then combined with the xy spline to generate a xyz spline, which is then used to generate a straightened 3D volume (②). The straightened 3D volume is finally projected on the xy plane and its 1D intensity profile Is extracted by averaging intensity values over the y axis. (③) for subsequent automated 1D autocorrelation analysis (④, see section below for details).

#### Autocorrelation analysis of sarcomere periodicity

Except for analysis reported in Figure 5D-G, autocorrelation analysis was performed using an automated procedure in Matlab (version 2019b) to detect periodicity in 1D titin intensity profiles (extracted from 1D ROIs, see previous section). The position and the prominence (amplitude) of the second correlation peak provides information about the spatial periodicity and the period regularity of the repetitive sarcomeric pattern (Figure 4D, panel ④).

For global autocorrelation analysis (Figure 4E-F’), the autocorrelation function was calculated over the total length of the 1D profiles and averaged by culture day (Figure 4E) or by morphological categories (Figure 4F). For local autocorrelation analysis (Figure 5), the autocorrelation function was locally applied on 10 µm wide (∼ 5 sarcomeres) sliding windows with a step size of 0.083 µm along the 1D profile. For each sliding window, the position and the prominence of the second autocorrelation peak, were automatically extracted and the local portion of the fiber was classified as periodic if the prominence exceeded 0.6 (Figure 5A). Regions shorter than 6 µm were discarded as noise. Neighbouring regions less than 10 µm apart were connected. Four further quantities were calculated for each 1D titin intensity profiles: the proportion of periodic regions (Figure 5B); the average length of consecutive periodic regions (Figure 5C), the number of periodic regions per 100 µm of fiber (Figure 5D) 4) and the homogeneity score calculated as the standard deviation of periodic window positions (Figure 5E). Autocorrelation analysis of 1D titin intensity profiles was performed in Matlab (version2019b) with custom scripts. Autocorrelation analysis detects periodicity from temporal or spatial series, and displays the period as the distance of the secondary peak from the y axis (Figure 4D, panel ④).

Autocorrelation analysis reported in Figure 5D-G was performed in R using the acf() function from the “stats” package on manually segmented 1D intensity profile of live ɑ-actinin-mKate2 signals at several time points matched by myofibril morphology. The prominence of the 1^st^ ACF peak is automatically extracted from the resulting ACF profile (Figure 5E, G).

#### Quantifying and modelling recoils of laser ablated myofibers

Pairs of post-ablation severed ends were tracked manually for a duration of 5 seconds from the point of ablation onwards, with the Manual Tracking plugin in FIJI (Fabrice P. Cordelières, Institute Curie) and subsequently analysed in R with custom scripts. Mean and standard deviation of distances between pairs of severed ends were plotted against time (Figure 6B, dots and shaded areas, respectively).

Two types of models were fitted to each recoil time course in R: First, a linear model was fitted on recoils during the first second. The initial recoil velocity was extracted as the slope of the linear model (Figure 6B, dashed line and C). Second, a viscoelastic model was fitted using an asymptotic regression model from the drc package (Ritz et al., 2015) with the following equation:

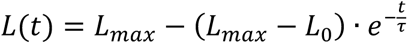

where the recoil distance L is a function of time t, L_max_ is the maximum attainable L, L_0_ is L at time = 0, and τ (viscoelastic relaxation time) is inversely proportional to the relative rate of L increase while t increases (Figure 6B, solid curves and D). τ is defined by the ratio of the material viscosity to the Young’s elastic modulus.

#### Statistical analysis and data visualization

Multiple pair-wise statistical tests in Figures 6 and 7 were performed using the paired.t.test() function in R. Normality of datasets were verified with the Shapiro test using the shaprio.test() function. Statistical significance was reported as p < 0.05. With the exception to Figure 4D-F’ and Figure 5A, all line and box plots were generated in R with standard plotting functions. Violin plots in Figure 5 were generated with the “vioplot” package (Adler and Kelly). Figure 4D-F’and Figure 5A were produced with Matlab, with Figure 4E’, F’ and Figure 5A generated with the Pseudocolor plot (pcolor) function.

## Acknowledgements

We thank all members of the Schnorrer group, the Pourquié group, the IBDM and the Centuri community for helpful discussions and feedback. We thank C. E. Seidman (Harvard University) and J. G. Seidman (Harvard University) for the gift of GFP-titin iPSC line, B. Friedrich (TU Dresden) for valuable inputs on algorithms for vector field analysis, the IBDM imaging facility for microscopy support, and D. Massey-Harroche (IBDM), E. Bazellières (IBDM), V. Dupuis (IBDM) and S. Tlili (IBDM) for cell culture support.

## Supplemental figure legends

**Figure 1 – figure supplement 1.**
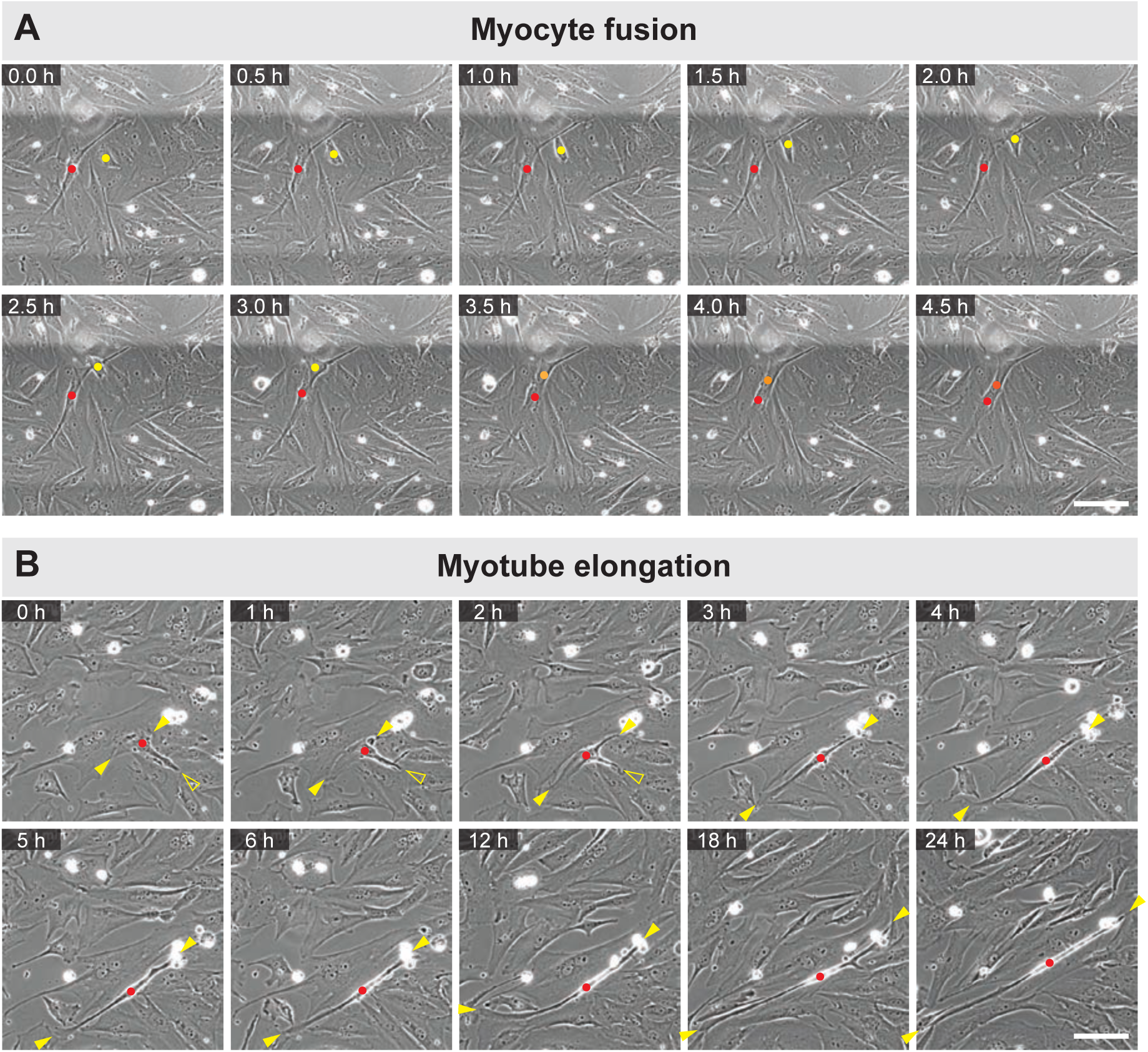
Dynamics of myocyte fusion and myotube elongation. Stills from phase contrast time-lapse movies of live myocultures. **A)** A myocyte (yellow dot) fusing into an elongated myotube (red dot). Fusion is highlighted by the orange-coloured dot of the newly fused nucleus. **B)** A single myotube (red dot) elongating. Yellow solid arrowheads show elongating ends, while yellow empty arrowhead shows a retracting extension. Scale bars: 100 µm. **[Related to Figure 1 and Figure 1 – video 1.]**

**Figure 2 – figure supplement 1.**
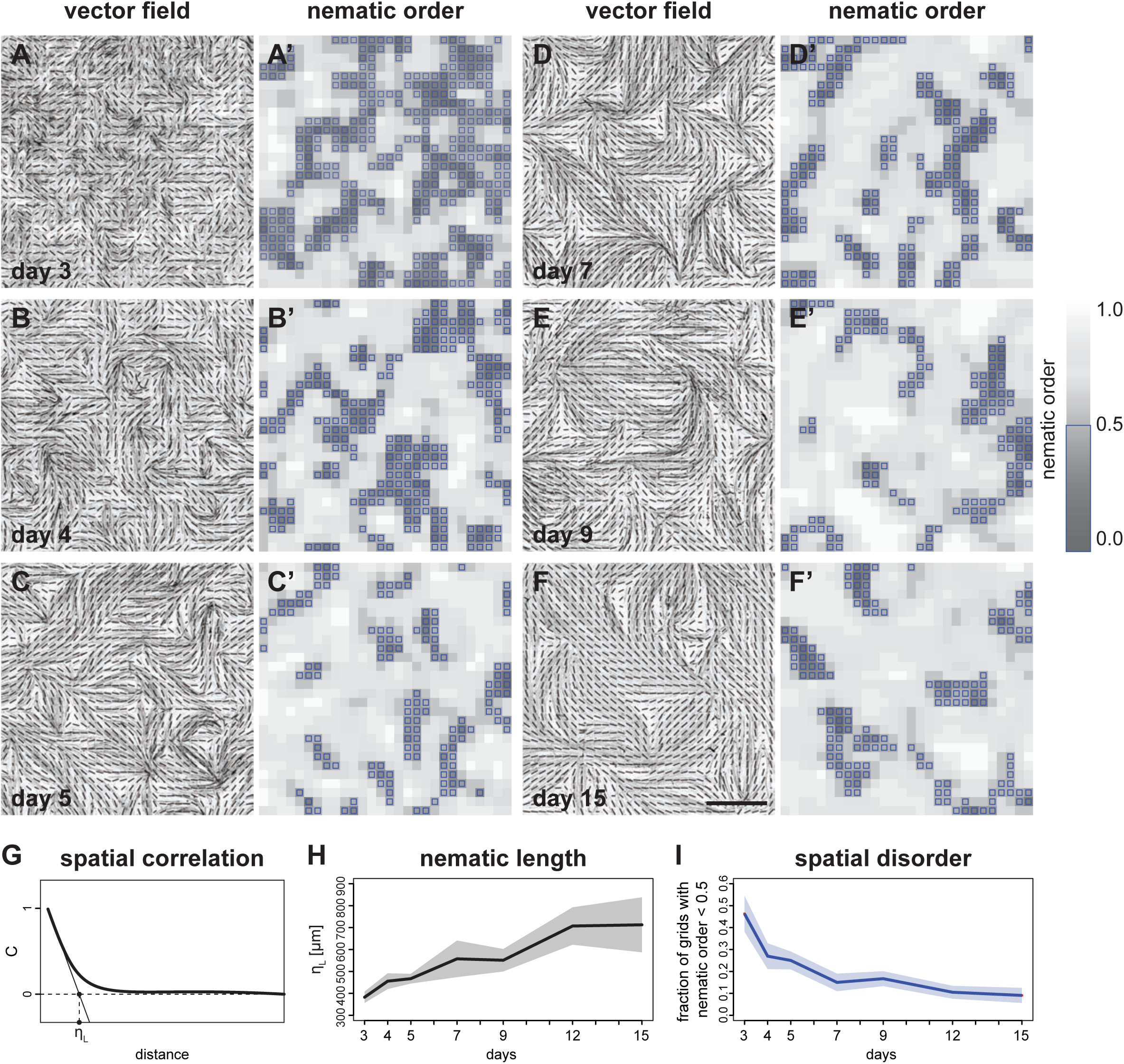
Vector field analysis of myofiber bundle alignment. **A-F)** inverted grey scale images of phalloidin staining superimposed with grids of vector fields generated from OrientationJ. Vector orientation and coherence are represented by the direction and length of each short line. **A’-F’)** 2D heatmaps showing local nematic order *S* for each grid position. Dark regions show misalignment (lower values), while light regions show good alignment (higher values). Blue boxes highlight grids with nematic order <0.5. Colour map shown on the right. **G**) A hypothetical spatial autocorrelation function from a vector field. η_L_ is the nematic correlation length. **H)** Time course of the nematic correlation length η_L_ (n = 6 windows per stage). **I)** Time course of the fraction of grids with local nematic order *S* < 0.5 (n = 6 windows per stage). In H and I, solid lines represent the mean, while shaded areas show standard deviation. Scale bar: 1mm. **[Related to Figure 2. Source data available for panels H and I.]**

**Figure 2 – figure supplement 2.**
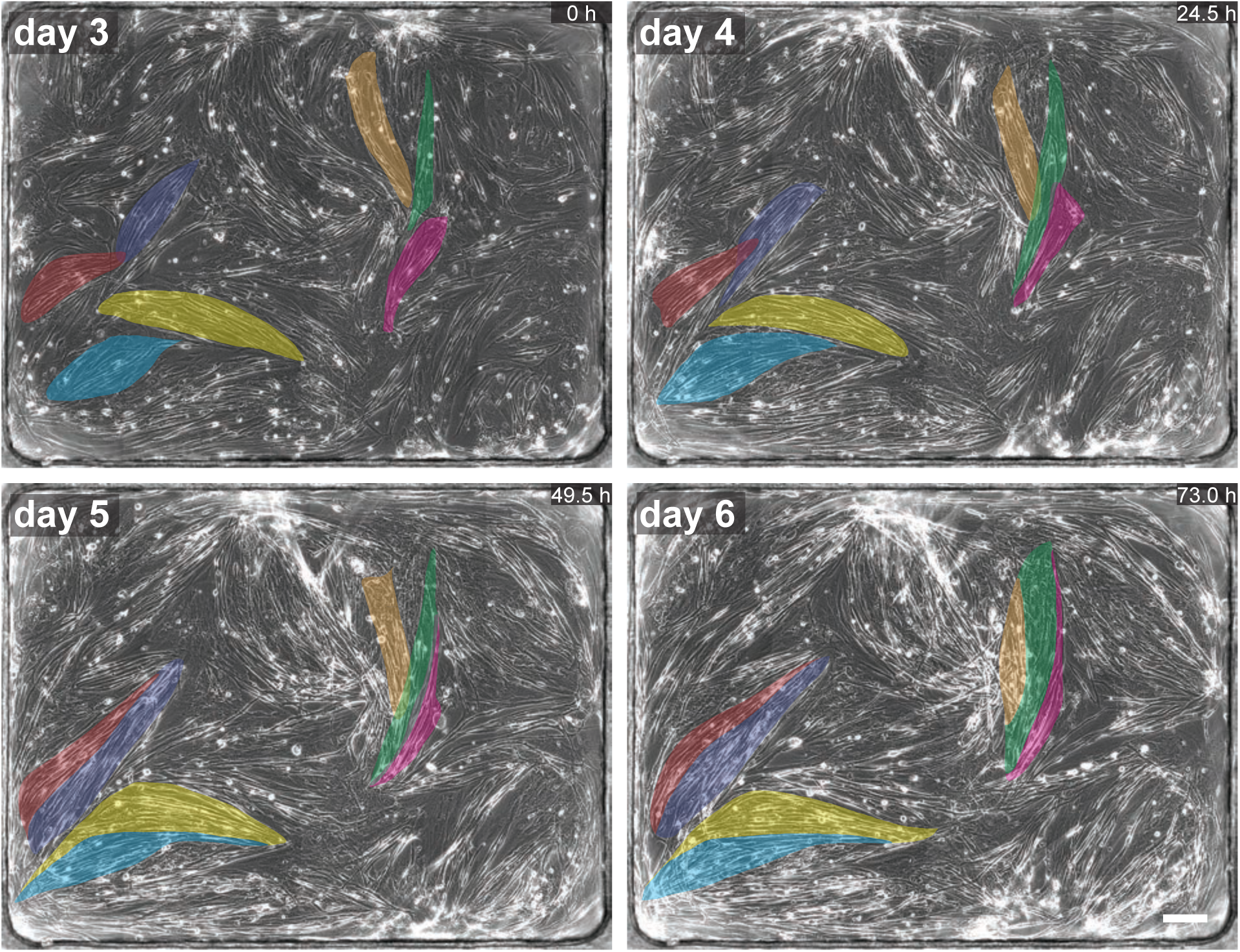
Dynamics of myofiber bundles. Phase-contrast imaging of myocultures showing myofiber bundles merging during the first 6 days of differentiation. Colour-shaded areas highlight individual myofiber bundles. In each of the three events highlighted, two or three myofiber bundles merge into one. Scale bar: 200 µm. **[Related to Figure 2 and Figure 2 – video 1.]**

**Figure 3 – figure supplement 1.**
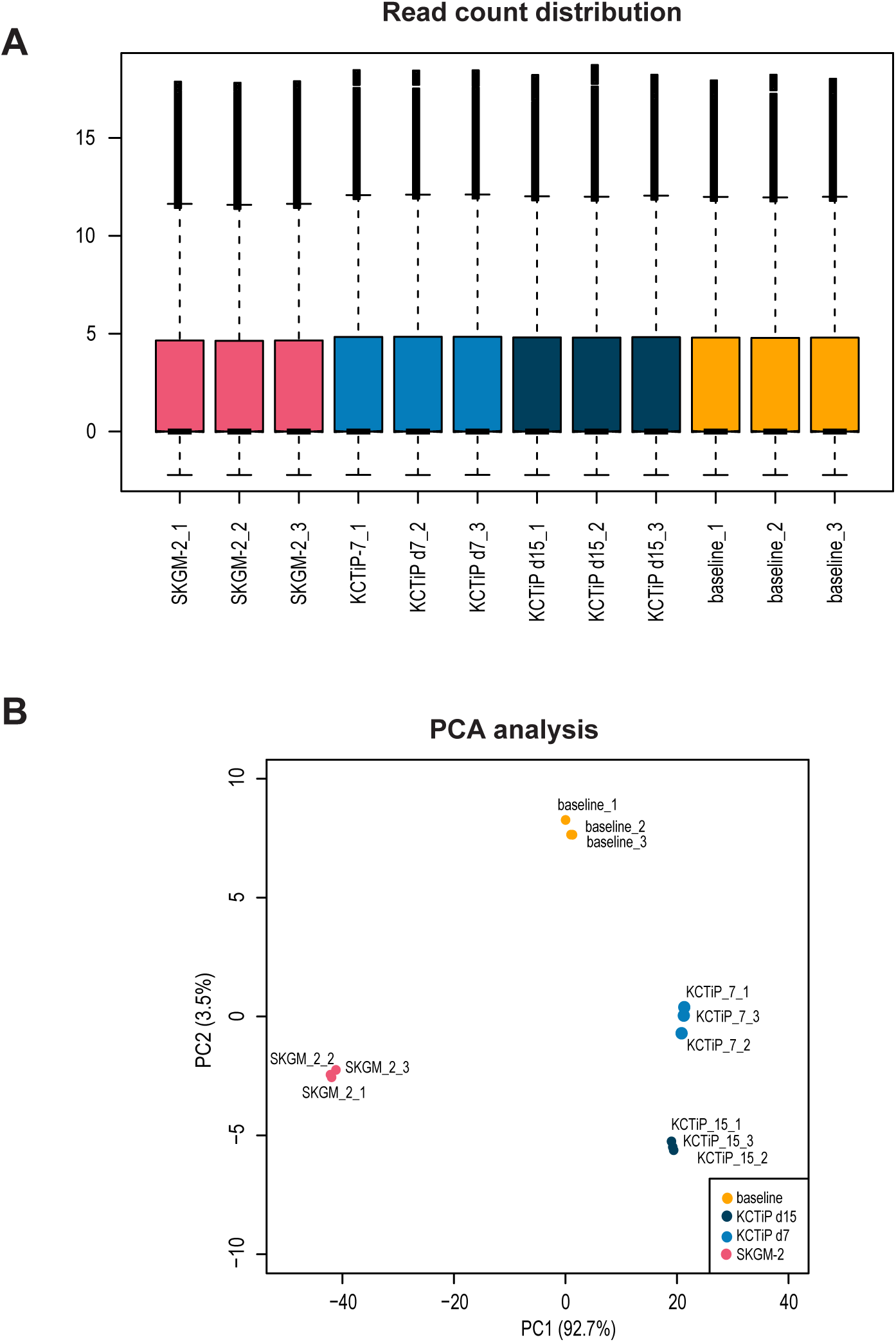
Quality control of the transcriptomics data set. **A)** Read count distribution for all biological replicated analysed. **B)** Principal components analysis shows the strong similarity of the replicates.

**Figure 5 – figure supplement 1.**
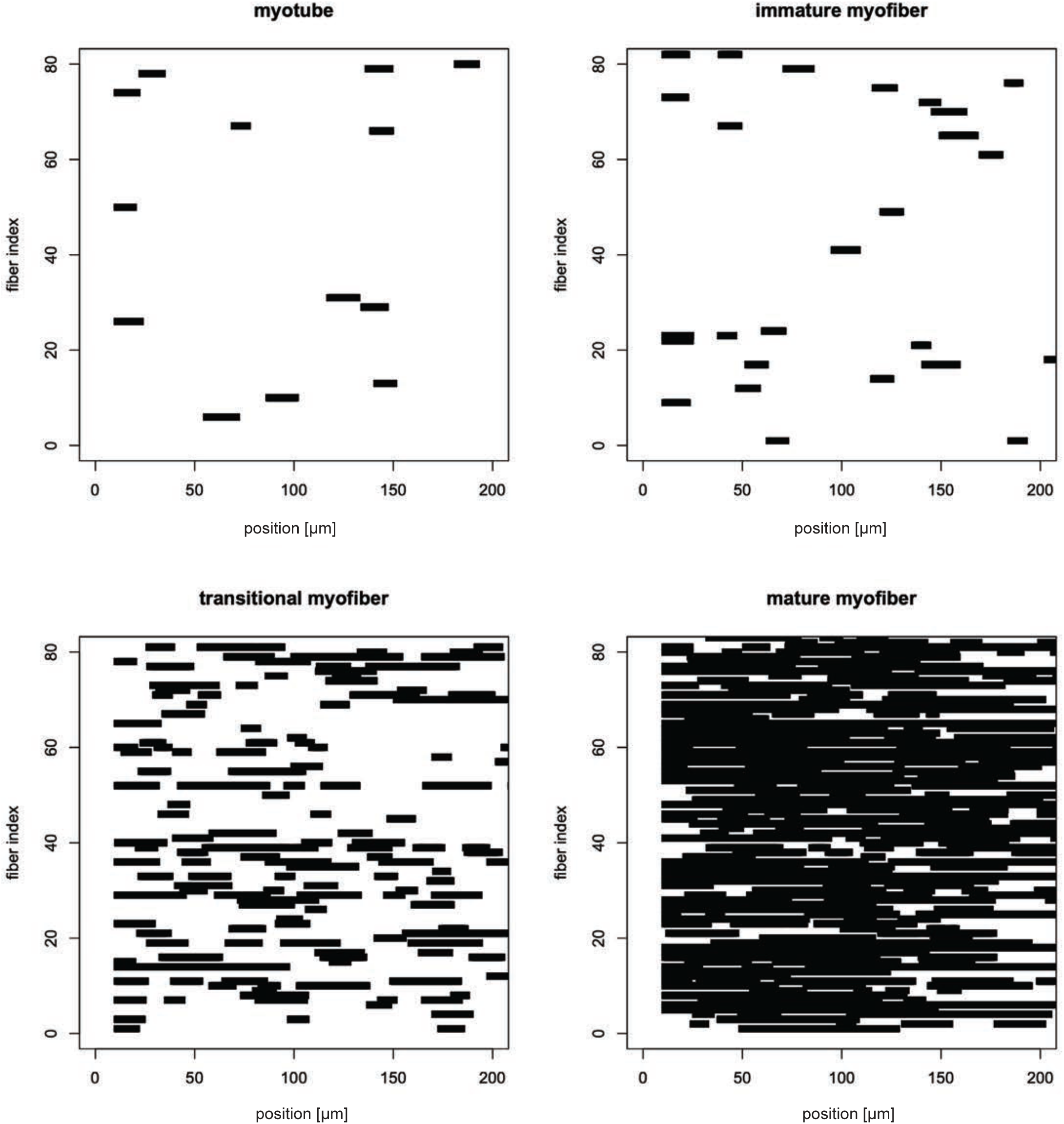
Distribution of patterned regions in randomly selected myotubes, immature, transitional and mature myofibers. Distribution of patterned regions (ACF peak prominence >0.6, see methods) in randomly selected myotubes, immature, transitional and mature myofibers. In each plot, the x axis displays positions along the myofiber in µm, while each myofiber occupies a row along the y axis. Note the homogenous distribution of the pattern regions. **[Related to Figure 5.]**

**Figure 6 – figure supplement 1.**
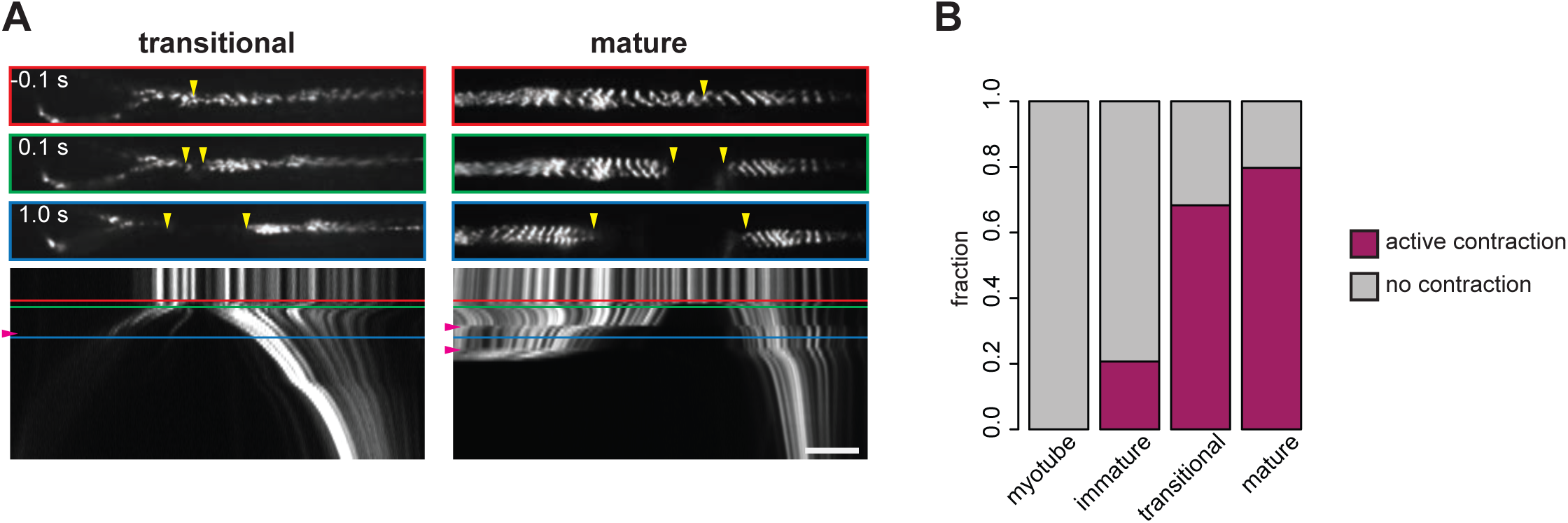
Contraction of myofibers revealed by point laser ablation. **A)** Examples of contracting transitional (left) and mature myofibers (right) post laser ablation. Top, upper middle and lower middle panels show 0.1 s prior to, 0.1s after and 1.0 after ablation, respectively. Bottom panels show the kymographs. Coloured boxes correspond to coloured lines in the kymograph. Yellow vertical arrowheads show sites of ablation and severed ends. Purple horizontal arrowheads on kymographs highlight jerky motions caused by contraction events. **B)** Frequency of contraction events in myotubes, immature, transitional and mature myofibers. Scale bar: 100 µm. **[Related to Figure 6 and Figure 6 – video 2. Source data available for panel B.]**

**Figure 6 – figure supplement 2.**
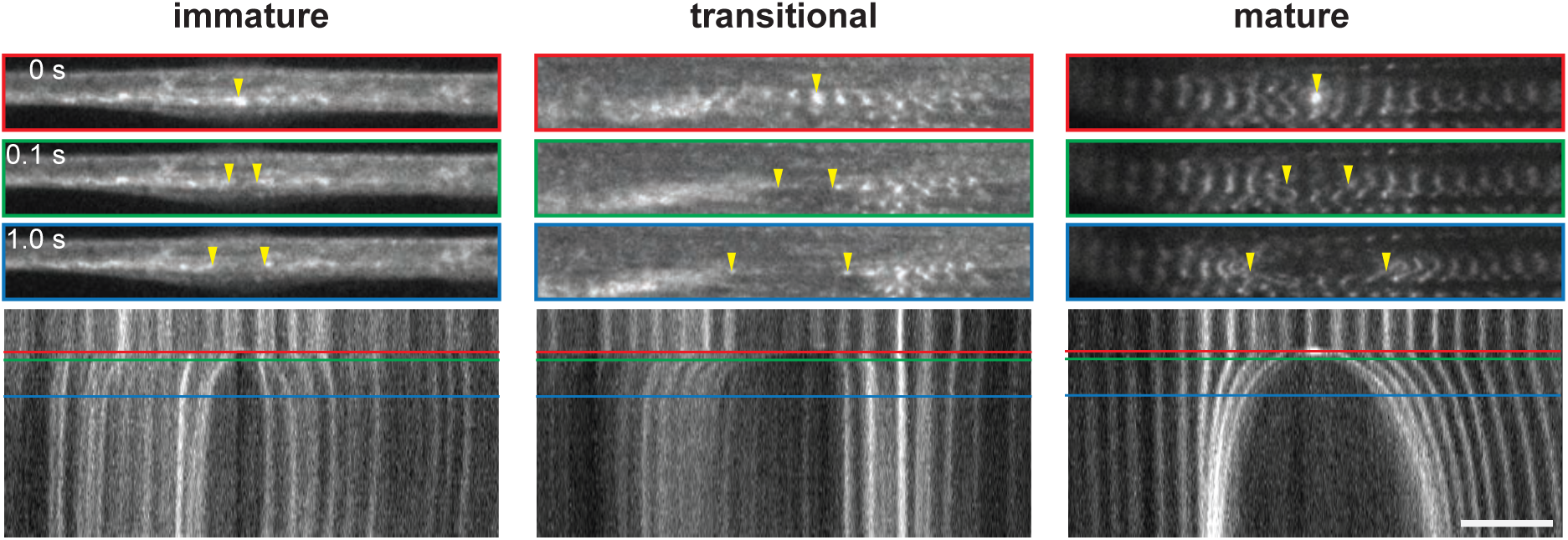
Tension on myofibrils revealed by point laser ablation. Recoil of myofibrils in an immature (left), a transitional (middle) and a mature myofiber (right) post point laser ablation. Top, upper middle and lower middle panels show the ablation events, 0.1s after and 1.0 after ablation, respectively. Bottom panels show the kymographs. Coloured boxes correspond to coloured lines in the kymograph. Yellow vertical arrowheads show sites of ablation and severed ends. Note that the mature sarcomeres collapse after the cut. Scale bar: 100 µm. **[Related to Figure 6 and Figure 6 – video 3.]**

**Figure 7 – figure supplement 1.**
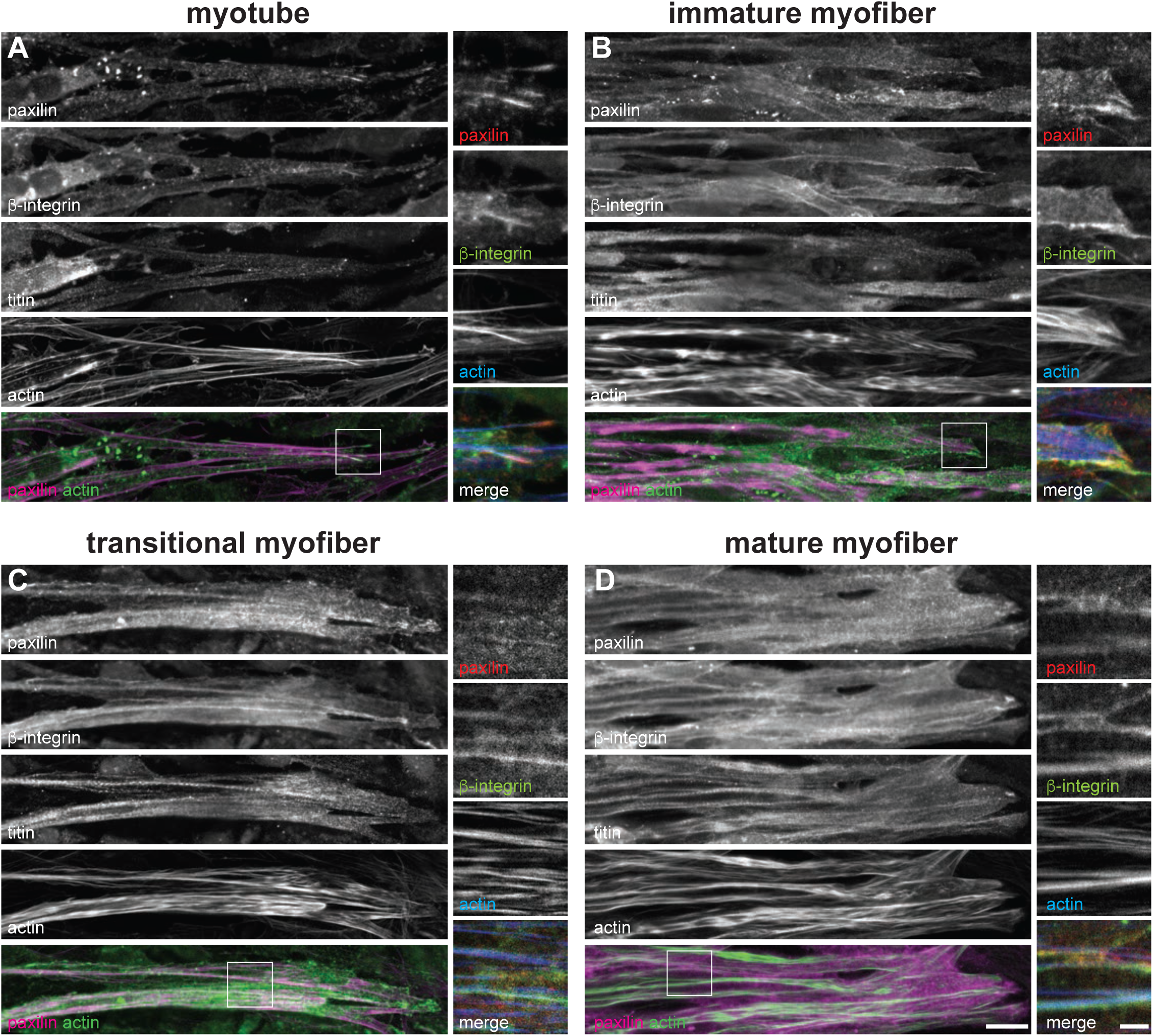
Maturation of myofiber attachments revealed by additional attachment markers. **A-D)** Immunostainings of paxillin (top), β1-integrin (upper middle), titin (middle), actin (lower middle) and a merge of paxillin and actin signals (bottom) in myotubes (A), immature (B), transitional (C) and mature myofibers (D). Insets shows a larger zoom on the right, with paxillin (top), β1-integrin (upper middle), actin (lower middle) and a merge of paxillin, β1-integrin and actin signals (bottom). Scale bars: 20 µm in smaller zoom and 5 µm in larger zoom.

**Figure 7 – figure supplement 2.**
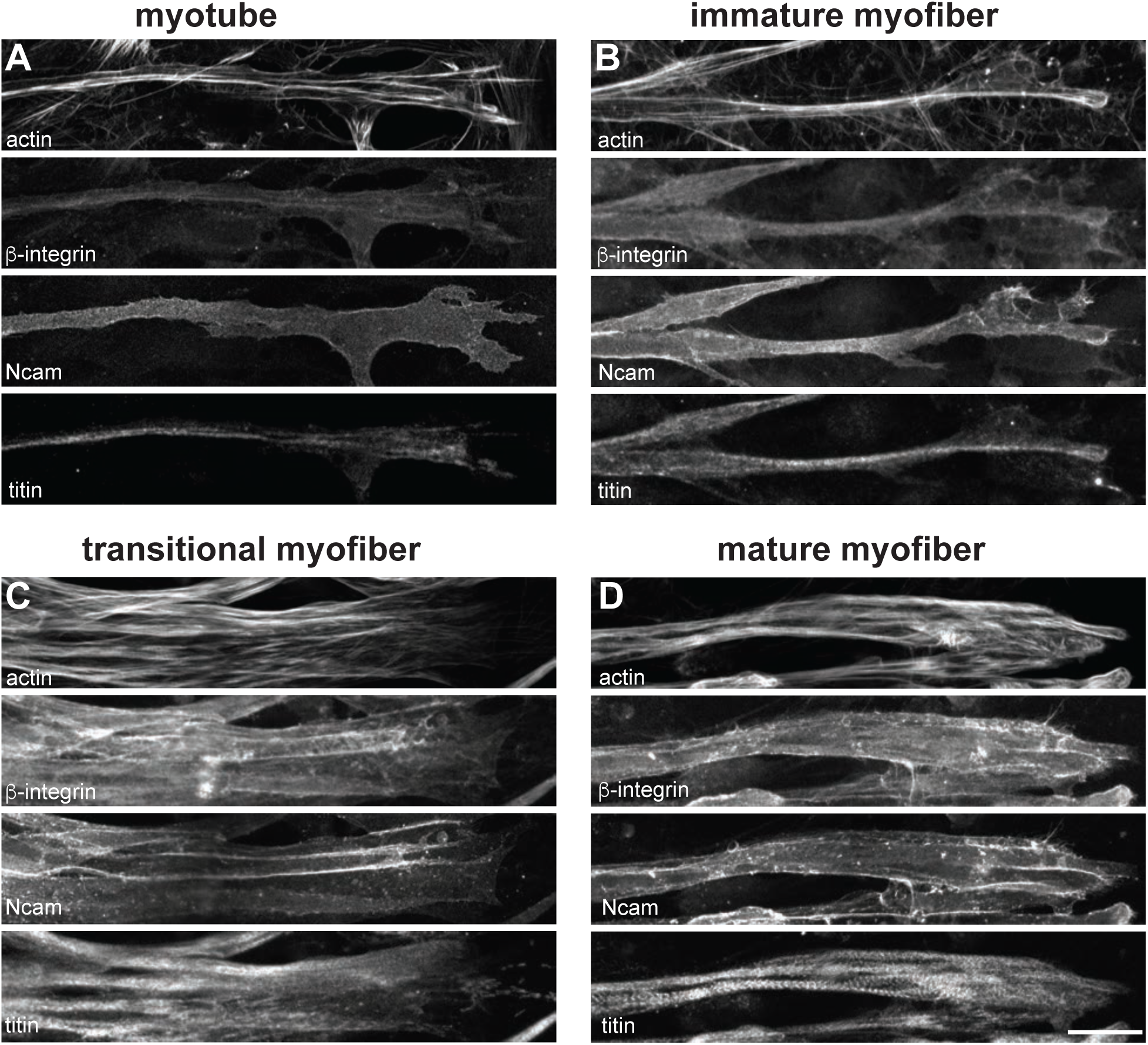
Maturation of myofiber attachments revealed by additional attachment markers. **A-D)** Immunostainings of actin (top), β1-integrin (upper middle), NCAM1 (lower middle) and titin (bottom) in myotubes (A), immature (B), transitional (C) and mature myofibers (D). Scale bars: 20 µm. **[Related to Figure 7.]**

## Movie legends

**Figure 1 – video 1. Phase contrast movie of live myoculture showing a myocyte fusing into an already elongated myotube, and the elongation of a myotube**

Time resolution: 30 min per frame. Duration: 4.5 hours and 24 hours, respectively. Scale bar: 100 µm. **[Related to Figure 1 – figure supplement 1.]**

**Figure 2 – video 1. Phase contrast movies of a live myoculture showing two myofiber bundles merging**

Upper: original images superimposed with highlighted myofiber bundles. Lower: processed images pseudo-coloured in OrientationJ. Time resolution: 30 min/frame. Duration: 74 hours. Scale bar: 200 µm. **[Related to Figure 2 – figure supplement 2.]**

**Figure 5 – video 1. A time-lapse movie showing a myofiber labelled with mKate2-α-actinin2undergoing myofibrillogenesis**

Time resolution: 15 min/frame. Duration: 47.75h. Scale bar: 20 µm**. [Related to Figure 5.]**

**Figure 5 – video 2. A time-lapse movie showing end retraction in a myofiber labelled with α-mKate2-α-actinin2**

Time resolution: 15 min/frame. Duration: 4h. Scale bar: 20 µm**. [Related to Figure 5.]**

**Figure 6 – video 1. Line laser ablation experiments for myotube, immature, transitional and mature myofibers.**

Time resolution: 100 ms/frame. Duration: 6 seconds. Scale bar: 100 µm. **[Related to Figure 6.]**

**Figure 6 – video 2. Contraction of myofibers during laser ablation**

Time resolution: 100 ms/frame. Duration: 6 seconds. Scale bar: 100 µm. **[Related to Figure 6 – figure supplement 1.]**

**Figure 6 – video 3. Recoil and collapse of myofibrils post point laser ablation**

Time resolution: 100 ms/frame. Duration: 6 seconds. Scale bar: 100 µm. **[Related to Figure 6 – figure supplement 2.]**

## Table legends

**Supplementary Table 1. Raw and normalised RNAseq data of undifferentiated (SKGM), day 7 and day 15 myogenic cultures. GO-term enrichments of all 10 Mfuzz clusters**

